# Association analyses reveal both anthropic and environmental selective events during eggplant domestication

**DOI:** 10.1101/2024.11.25.625206

**Authors:** Emmanuel Omondi, Lorenzo Barchi, Luciana Gaccione, Ezio Portis, Laura Toppino, Maria Rosaria Tassone, David Alonso, Jaime Prohens, Giuseppe Leonardo Rotino, Roland Schafleitner, Maarten van Zonneveld, Giovanni Giuliano

## Abstract

Eggplant (*Solanum melongena*) is one of the four most important Solanaceous crops, widely cultivated and consumed in Asia, the Mediterranean basin, and Southeast Europe. We studied the Genome-wide association (GWA) of historical genebank phenotypic data on a genotyped worldwide collection of 3,449 eggplant accessions. Overall, 334 significant associations for key agronomic traits were detected. Significant correlations were obtained between different types of phenotypic data, some of which were not obvious, such as between fruit size/yield and fruit color components, suggesting simultaneous anthropic selection for genetically unrelated traits. Anthropic selection of traits like leaf prickles, fruit color, and yield, acted on distinct genomic regions in the two domestication centers (India and Southeast Asia), further confirming the multiple domestication of eggplant. To discriminate anthropic from environmental selection in domestication centers, we conducted a Genotype-Environment Association (GEA) on a subset of georeferenced accessions from the Indian subcontinent. The population structure in this area revealed four genetic clusters, corresponding to a latitudinal gradient, and environmental factors explained 31% of the population structure when the effect of spatial distances was removed. GEA and outlier association (OA) identified 305 candidate regions under environmental selection, containing genes for abiotic stress responses, plant development, and flowering transition. Finally, in the Indian domestication center anthropic and environmental selection acted largely independently, and on different genomic regions. These data allow a better understanding of the different effects of environmental and anthropic selection during domestication of a crop, and the different world regions where some traits were initially selected by humans.

## Introduction

Eggplant (*Solanum melongena* L.) is, with potato, tomato, and pepper, one of the four most cultivated Solanaceous food crops, with a worldwide production exceeding 58.6 M tons in 2021 (FAO, 2022). About 17,700 eggplant accessions are currently listed in the Genesys worldwide catalog (https://www.genesys-pgr.org/) and maintained by dozens of genebanks. Eggplant has extensive genetic diversity adapted to different environments (Solberg *et al*., 2023), so its genome is a potential reservoir of adaptive traits. Discovering the genes controlling these traits is crucial for the future improvement of this crop.

As genebanks evolve from germplasm providers to genetic resource knowledge centers, new opportunities emerge for understanding crop diversity (CGIAR, 2020). Genebanks are invaluable providers of phenotypic data for highly heritable traits, helping users to select germplasm and understand trends and patterns in trait evolution of crop genepools. Therefore, genebank genomics represents an important tool for understanding the genetic structure of such genepools. In fact, besides supporting the identification of duplicate samples within collections, and the correction of taxonomic misassignments, genomic tools can also be used for Genome-wide association (GWA) studies on historical phenotypic data collected by genebanks during seed multiplication (Tripodi *et al*., 2021; Langridge and Waugh, 2019; Milner *et al*., 2019). Among the technologies used for genotyping, the Single Primer Enrichment Technology (SPET) (Barchi *et al*., 2019; Scaglione *et al*., 2019; Herrero *et al*., 2020; Villanueva *et al*., 2021) represents a customizable, cost-efficient solution although it requires *a priori* genomic or transcriptomic information and identification of SNPs for probe design. Recently, a large worldwide collection of 3,499 eggplant accessions, including landraces and wild relatives, was characterized using SPET, identifying its population structure, centers of origin, and migration routes (Barchi *et al*., 2023).

The genetic diversity safeguarded in genebank collections can be used to overcome challenges posed by climate change, such as rising temperatures and unpredictable rainfall patterns, which are increasingly likely to affect crop productivity. Assessing phenotypic resilience in field conditions remains laborious and costly, so there is a pressing need for new approaches for screening large germplasm collections for adaptive traits. Genome-environment association (GEA) methods that link genetic information with environmental factors (Manel *et al*., 2010) are an example of such alternative approaches. Novel statistical methods and the growing accessibility of global-scale environmental data are opening the possibility of estimating the influence of the environment on crop genomic variation. A frequently applied GEA method is redundancy analysis (RDA), a multivariate statistical method based on expanding multiple regression for analyzing data with multivariate response data (Legendre and Legendre, 2012). Recent studies have shown that GEA can identify potential adaptive loci and populations that are adapted to stressful conditions in wheat, sorghum, barley and African wild eggplants (Chang *et al*., 2022; Lei *et al*., 2019; Lasky *et al*., 2015; Costa-Neto *et al*., 2023; Omondi *et al*., 2024).

In this study, we performed GWAS for 23 agronomical traits on a previously genotyped worldwide eggplant collection, exploiting historical phenotypical data collected by genebanks to identify marker-trait associations for key traits under anthropic selection. Additionally, we used a subset of 324 georeferenced accessions from one of the eggplant’s proposed centers of domestication, the Indian subcontinent, for GEA studies. This study aims to identify and compare genomic regions under anthropic and environmental selection, to identify the key environmental variables with the largest genomic influence, and to identify which traits selected by humans in the domestication centers spread to the rest of the world.

## Results and Discussion

### GWA analysis of historical genebank phenotypic data reveals regions under human selection in an eggplant worldwide collection

Agro-morphological data about traits routinely phenotyped by genebanks during seed multiplication and characterization trials, were collected for a previously genotyped worldwide eggplant collection comprising 3,499 accessions (Barchi *et al*., 2023) and manually curated to be comparable (**Table S1**). A total of 23 phenotypic traits were checked for consistency, resulting in sets of observations ranging from 438 for fruit size (frusiz) to 1,517 for fruit color (frucol) (**Table S2, S3**).

Several phenotypic traits displayed a biased geographical distribution. For instance, prickly eggplant types are preferred in the two putative domestication centers identified by Barchi *et al*. (2023) (India and Southeast Asia) based on their perceived superior organoleptic quality (**Figure 1a-b**), as are also anthocyaninless fruit types (**Figure 1c)**. Notably, these phenotypes resemble those of the eggplant direct progenitor, *S. insanum* (**Figure 1d**), a possible indication that these preferences reflect a cultural heritage dating back to early post-domestication.

**Figure 1:**
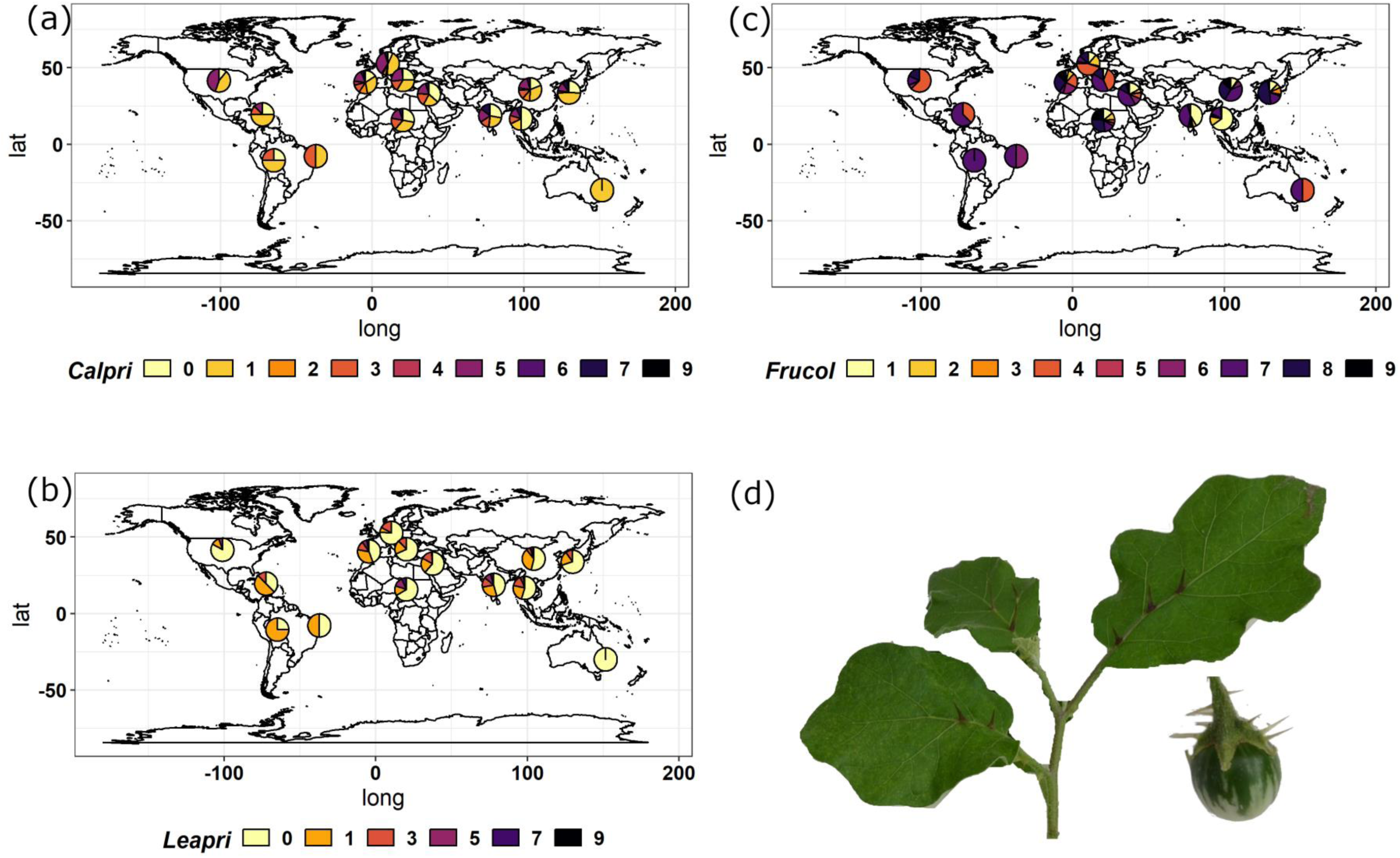
Geographical distribution of different eggplant traits: calyx prickles (a), leaf prickles (b) and overall fruit color (c). Color scales range from 0 (no prickles) to 9 (many prickles) and from 0 (no anthocyanins) to 9 (black fruits) (**Table S2**). (d) Leaf and fruit phenotypes of the eggplant progenitor*, S. insanum*.

Significant positive and negative inter-trait correlations were detected (**Figure S1**). Some of these correlations were expected: for example, fruit shape ratio (frushr) was positively correlated with fruit curvature (frucur) and negatively correlated with fruit section and fruit size (frusec and frusiz); also, leaf and calyx prickliness (leapri and calpri) were positively correlated. Other correlations were less obvious: fruit size (frusiz) and yield (fruyld) showed positive correlations with fruit color component frucol_a and negative correlations with frucol_b and frucol_L, probably because of a simultaneous selection for both fruit size and peel color in specific geographic areas. Additionally, growth habit showed small but significant negative correlations with fruit size and yield, with prostrate plants exhibiting smaller fruits and lower yields. This contrasts with what is observed in tomato F_2_ populations, where prostrate plants display higher yields, probably due to reduced lodging (Ozminkowski *et al*., 1990). Given the importance of yield as an agronomic trait, the genetic basis of this correlation merits further study.

To perform GWAS, the MLMM (Segura *et al*., 2012), FarmCPU (Xiaolei Liu *et al*., 2016), and BLINK (Huang *et al*., 2019) methods implemented in GAPIT3 (Wang and Zhang, 2021, p.3). Overall, 334 unique significant QTNs (p-value <= 0.01) were identified using the three methods (**Table S4**), corresponding to 229 non-redundant QTNs and 189 unique QTLs, assuming an LD decay of 100 Kbp. We detected several notable marker-trait associations for all key traits under selection, many corresponding to known quantitative trait loci (QTLs) and/or falling near genes with documented functions (**Table S5)**.

Prickles are a distinctive trait of eggplant and its wild relatives. However, their presence is an unwanted feature in modern cultivars since they damage fruits during bulk packaging and transportation, and has therefore been removed by breeding (Daunay *et al*., 2001). Plant prickles are modified glandular trichomes, and their biogenesis is controlled by a *LOG* gene (Satterlee *et al*., 2024) as well as by transcription factors from the MYB, bHLH, WD40, WRKY, NAC, GRAS, and/or C2H2 zinc finger families (Wang *et al*., 2019). Genes encoding transcription factors putatively involved in trichome/prickle formation were spotted within the QTLs identified for leaf and calyx prickles, alongside homologs of the Arabidopsis *GLABRA1* (*GL1*) gene and of genes involved in secondary cell wall biogenesis, like the strong candidate gene *STRUBBELIG-RECEPTOR FAMILY 2* (*SRF2*) on ch10, already reported to mediate tissue morphogenesis, biosynthesis of cellulose, ROS and stress-related gene induction as well as ectopic lignin and callose accumulation (Chaudary et al., 2020). Furthermore, significant QTNs were also found on chr. 8, in the same region already highlighted for prickles-related QTLs by Portis et al. (2014), containing a cluster of expansin-like proteins, and on chr. 6, in the same region containing a selective sweep for prickliness (Barchi *et al*., 2021) and the *LOG* prickliness gene (Satterlee *et al*., 2024), confirming that strong human selection for presence/absence of prickles was exerted on these regions (**Figure 2a, Table S5)**.

**Figure 2:**
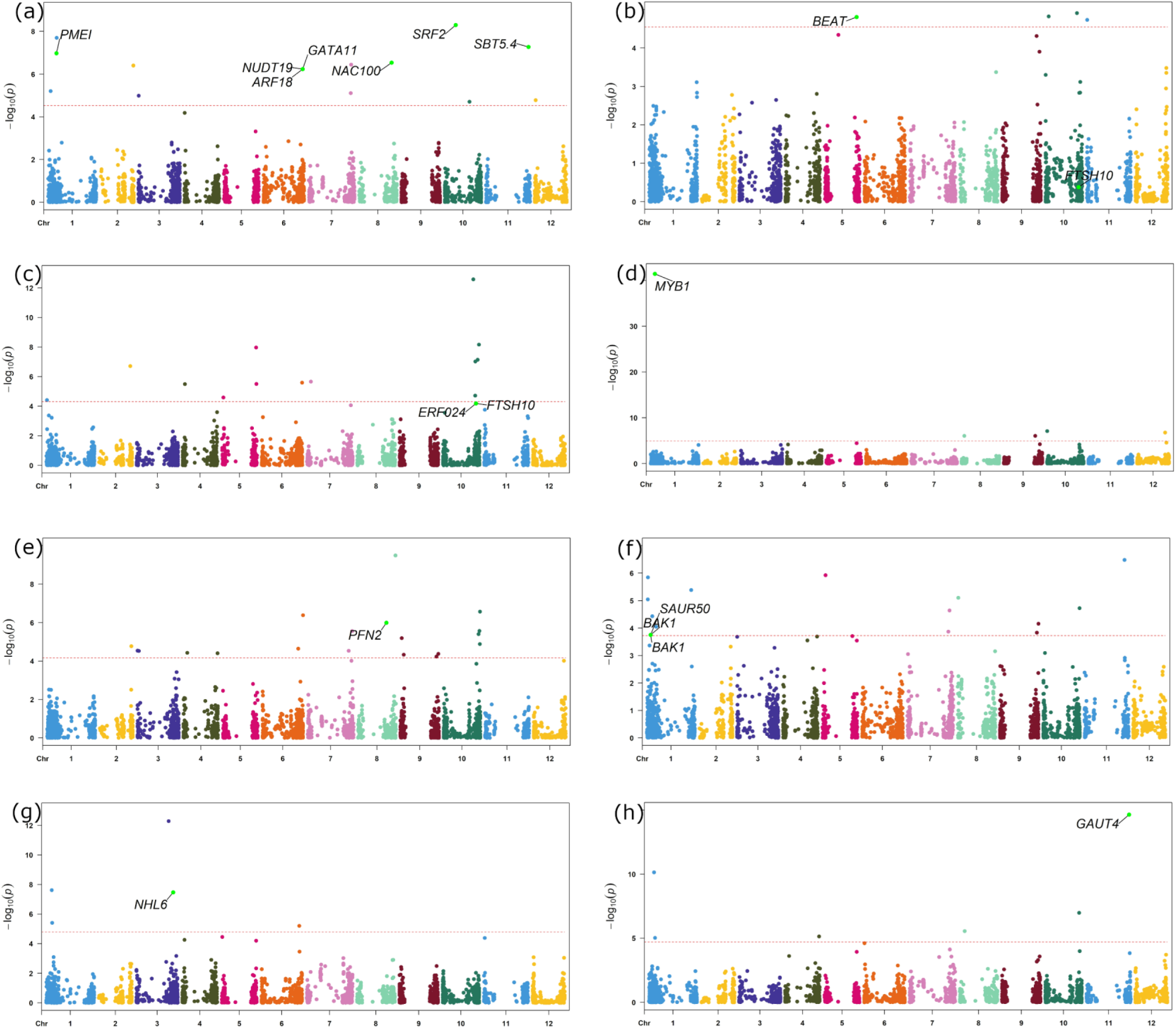
QTNs identified for leaf prickles (a); corolla color_b (b); fruit color_a (c); anthocyanin distribution (d); fruit cross section (e); fruit curvature (f), fruit apex (g) and growth habit (h). Red lines in the Manhattan plots indicate FDR significant level at p<0.01. Scale values, numbers and positions of significant QTNs are reported in Tables S1-S2. The most relevant candidate genes identified for the eight traits are also reported.

Anthocyanin pigmentation in eggplant is strongly dependent on the tissue, developmental stage, and environment (Zhang *et al*., 2014; Xiao *et al*., 2018; Barchi *et al*., 2019; Moglia *et al*., 2020). It is a distinctive trait of most modern eggplant varieties, and it allows ready identification of the vegetable in its various culinary forms. GWAS identified several significant QTNs for flower and fruit color, and anthocyanin accumulation in vegetative tissues. With a LD window of 100 Kb, several candidate genes were identified (**Figure 2b-c, Table S5).** In particular the transcription factors belonging to the MYB family (with *MYB1* as the most reliable candidate for the major QTN on Ch 01 with the highest P value), the homolog of phenylpropanoid biosynthetic genes *Anthocyanidin synthase* (*ANS*), plus an ATP-dependent zinc metalloprotease FTSH 10, putatively conferring non-photosensitivity of eggplant peel color (He *et al*., 2022) were identified by all the three models (BLINK, FarmCPU and MLMM), suggesting that these regions are deeply involved in the control of color/anthocyanin accumulation in eggplant. Additional QTLs, identified by one or two models, contain transcription factors belonging to ERF and BHLH families (Sun *et al*., 2016; Zhang *et al*., 2018), ABC transporters putatively mediating vacuolar transport of anthocyanins (Francisco *et al*., 2013) and the *MATE1* proanthocyanidin transporter (Pérez-Díaz *et al*., 2014). Other candidate genes include (i) the homolog of *Vitis vinifera Prx31*, involved in anthocyanin degradation (Movahed *et al*., 2016), (ii) the homolog of phenylpropanoid biosynthetic genes like *Flavone 3-dioxygenase* (*F3H-2*), (iii), and genes involved in the latest steps of anthocyanin decoration, like *Anthocyanidin 3-O-glucosyltransferase* (*3GT*) (Barchi *et al*., 2021), and the *SMEL_AAT* acyltransferase (Florio *et al*., 2021).

Several QTLs and QTNs control fruit flesh color. Among them, we identified several candidate genes associated with chlorophyll synthesis and photosynthetic complex biogenesis as potential candidates (**Table S5**). Furthermore, we also found significant QTNs on chr. 8, colocalizing with the *gring QTL* (Portis *et al*., 2014), as well as with *SmAPRR2*, recently identified to be associated with the absence of fruit chlorophyll pigmentation in a MAGIC population (Arrones *et al*., 2022) and with the *STAY-GREEN* gene (Sakuraba *et al*., 2015). Several genes and transcription factors belonging to the VOZ, GRAS, and TC families involved in the modulation of flowering time, photoperiod, and gibberellin signaling were identified within QTLs associated with controlling flowering time (**Table S5**).

GWAS analysis for fruit size/shape identified homologs of genes controlling these traits in tomato (**Figure 2d-e, Table S5)**, including *OVATE (*on chr. 1, within one of the major QTLs), *IQ-domain/SUN* (*IQD)*, and *CELL SIZE REGULATOR* (Mu *et al*., 2017). Interestingly, we also identified some candidate genes for fruit curvature (**Figure 2f, Table S5)**, including a *Growth-regulating factor* (*GRF*) gene involved in organ enlargement in Arabidopsis, rice and Citrus (Xiao Liu *et al*., 2016), a probable auxin efflux carrier component 1c (*PIN1C*), involved in fruit curvature in cucumber (Li *et al*., 2020) as well as a member of auxin-responsive *Small Auxin-Up RNA* (*SAUR*) genes (Stortenbeker and Bemer, 2019).

The *POINTED TIP* (*PT*) gene and its alleles, *PT^H^*and *PT^R^*, regulate the development of tomato fruit with or without pointed tips by differentially affecting downstream genes (Song *et al*., 2022). *PT* encodes a C2H2-type zinc finger protein. In Arabidopsis, the homologous *IDD15/SGR5*, *IDD14,* and *IDD16*, controls organ morphogenesis and gravitropism through auxin biosynthesis and transport. Unlike Arabidopsis, tomato *PT^H^* and *PT^R^* have three and two zinc finger domains, respectively. Phylogenetic analysis shows PT is distinct, with SE3.1 being its closest homolog in tomato, involved in style extension and self-fertilization. This implies PT’s unique role in tomato fruit morphology. Our analysis did not highlight any homolog of *PT* in the QTLs identified for fruit apex. However, a gene in the QTL on chromosome 7 (**Figure 2g, Table S5)** was annotated as *ARF5*, involved in the auxin signaling; although auxin was reported to be involved in the development of fruit with pointed tips, the mechanism is still largely unknown (Song *et al*., 2022).

Within the QTLs controlling growth habit, a *GAUT4* gene was identified in a QTL on chromosome 11 by all the GWA models. In tomato, *GAUT4*-silenced plants exhibited an increment in vegetative biomass associated with palisade parenchyma enlargement (de Godoy *et al*., 2013), as well as in taller plants. This may suggest the involvement of this gene in difference between upright and prostate eggplant accessions. In addition, some *SAUR* genes, known to be involved in the regulation of dynamic and adaptive growth, as well as in stress responses (Zhang *et al*., 2021), were mapped (**Figure 2h**, **Table S5)**, as also a homolog of the WALLS ARE THIN 1 (*WAT1*) gene, known in Arabidopsis to confer a show a shorter and bushier phenotype in *wat1* mutants compared to functional QAT1 plants (Ranocha *et al*., 2010), suggesting a putative role of this gene in the control of growth habit also in eggplant.

### Different traits/genomic regions were selected in the two eggplant domestication centers

Recently, Barchi et al. (2023) identified a clear separation of the genetic ancestry of accessions from India and Southeast Asia, suggesting the existence of separate domestication centers in these two regions. To investigate whether human selection acted on similar or different genomic regions during domestication in the two regions, separate Genome-Wide Association (GWA) studies on the examined 23 traits (**Table S6**) were conducted on accessions of India and Southeast Asian origin. These studies were therefore compared with results obtained from the complete worldwide collection, revealing the exclusive origin of some associations (**Figure 3**). In the overall analysis, leaf prickles were associated to a genomic region on chromosome 3 and 6: the former association was found only in the Southeast Asian while the latter only in accessions from India. Regarding calyx prickles, QTNs on chromosomes 5 and 8, identified across all panels, were also found in the Indian accessions. On the other side, a QTN on chromosome 8 (around 89 Mbp) found in the overall analysis was identified also in the Southeast Asian accessions, suggesting that human selection may have acted on different genomic regions based on geographical area.

**Figure 3:**
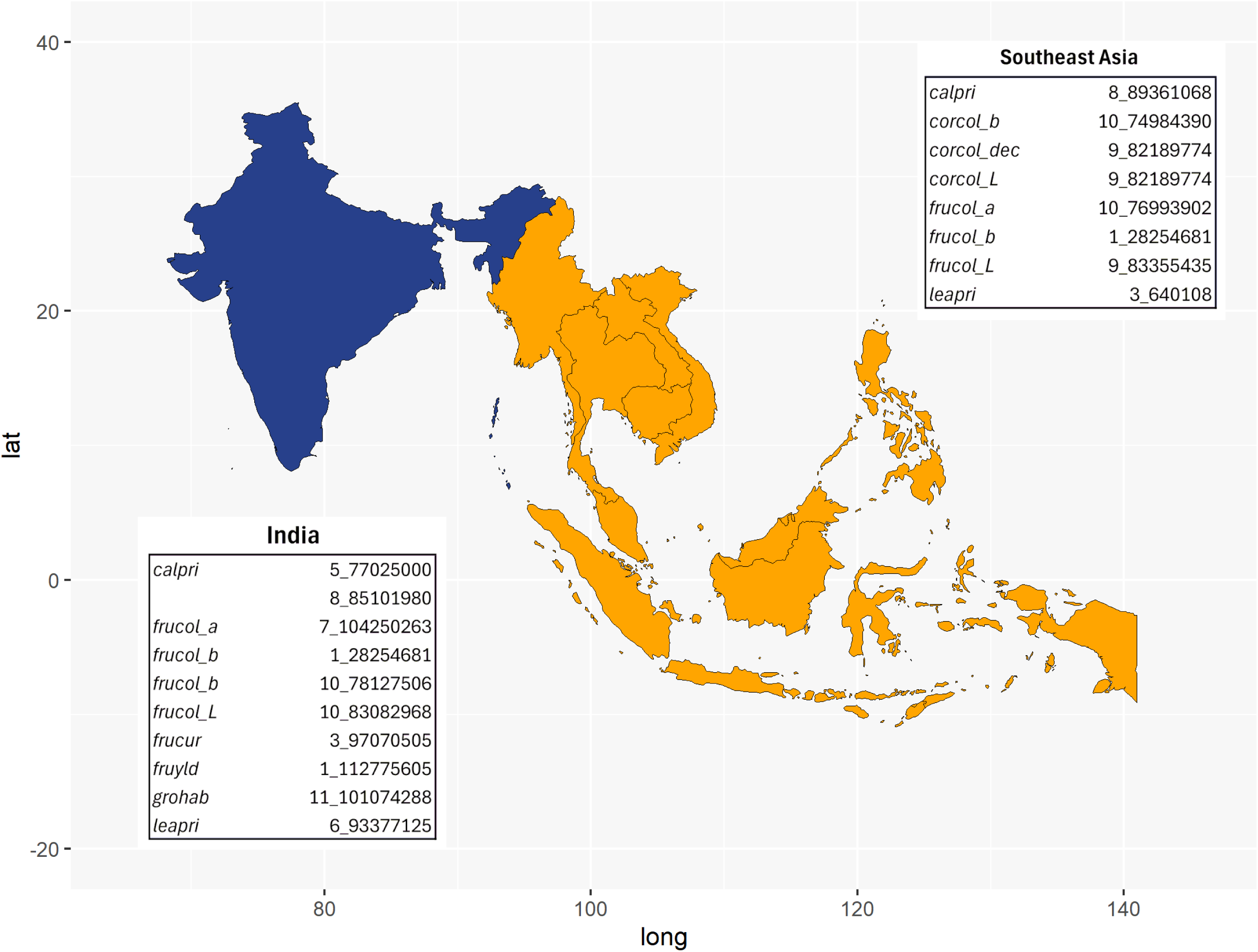
Different traits/ genomic regions were selected by humans in the two eggplant domestication centers. For each of the domestication centers, the traits/QTNs showing the strongest associations by GWA analysis are reported.

Interestingly, accessions exhibiting a pale corolla color (white or green) appear to have been primarily selected in Southeast Asia. In this region, two QTLs, on chromosome 9 (82.09 Mbp) and 10 (74.98 Mbp) were identified in both Southeast Asian and overall analyses. Notably, the former region contains an ABC transporter gene putatively involved in vacuolar transport of anthocyanins (Francisco *et al*., 2013). Furthermore, the selection for fruit color may also have occurred in common as well as different genomic regions for Southeast Asian and Indian accessions. Indeed, a common QTL was found for *frucol_b* on chromosome 1 at 28.25 Mbp. On the one hand, for Southeast Asian accessions, the human selection may have acted on chromosome 9 (83.35 Mbp) and 10 (76.99 Mbp). On the other hand, human selection for Indian accessions may have occurred within chromosome 7 (104.42 Mbp), were a BHLH120 was found, and in two genomic regions on 10 (78.12 and 83.08 Mbp), although no promising candidate genes were found.

Fruit yield, size, and shape are crucial agronomic traits targeted by humans during crop domestication. Our GWA analysis highlighted a region on chromosome 1 (112.6 Mbp) containing an ETHYLENE INSENSITIVE 3-like protein-coding gene that appears to have been selected specifically in India for fruit yield. Additionally, chromosome 3 (97.07 Mbp) harbors a QTL identified in both Indian and overall analyses, where IQ-domain/SUN (IQD) genes were found. Finally, a putative region under human selection in India is on chromosome 11 (100.9 Mbp) for growth habit within a QTL containing a *GAUT4* gene.

### Environmental selection in the Indian subcontinent

The Indian subcontinent accessions were subjected to Bayesian clustering using STRUCTURE. This analysis identified four clusters (K=4) (**Figure 4a**; **Figure S2**). The geographic distribution of the population structure, plotted as pie charts on the India-Bangladesh maps, clearly showed a directional distinction of the populations based on the grouping by regions (**Figure 4b)**. Accessions from Northeastern India and some populations from Eastern India cluster together with populations from Bangladesh. East and Northeast Indian regions comprised individuals modeled as highly admixed compared to the other areas. The Principal Component Analysis (PCA) clustering corresponded to the STRUCTURE results (**Figure 4c**). The first two PC axes explained 16.5% and 4.2% of the variation, respectively. The Indian populations separated from Bangladeshi populations on the first axis.

**Figure 4:**
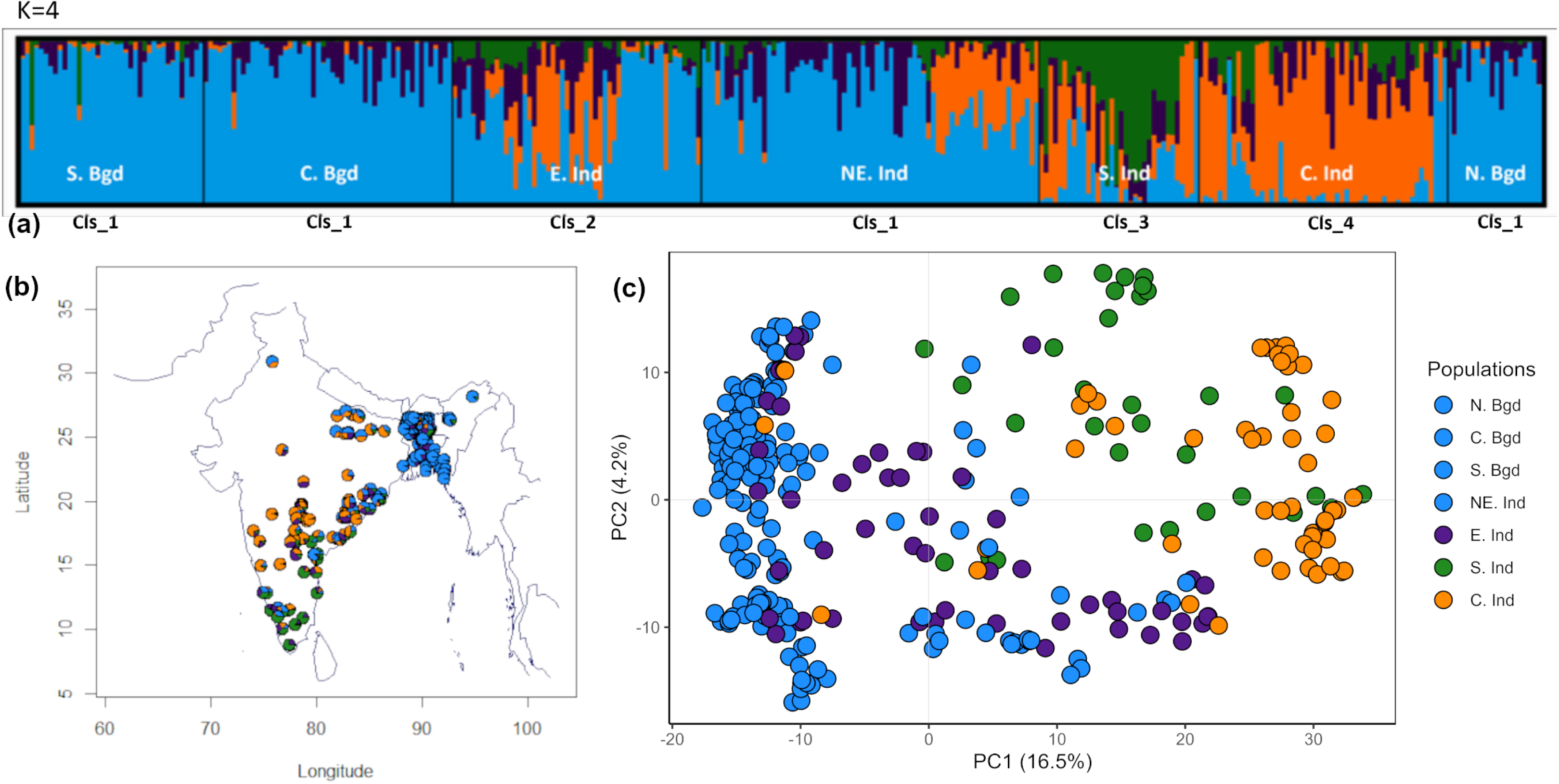
Genetic population structure of accessions from the Indian subcontinent. (a) Admixture proportions of the accessions as identified with STRUCTURE and identified by clusters 1-4 corresponding to the K=4 identified by K-means, (b) pie charts representing the ancestry coefficient of the eggplant accessions estimated using STRUCTURE; (c) Principal Component Analysis based on genomic data. The points are colored by cluster membership to correspond with the STRUCTURE results.

Among the ten environmental variables were retained for downstream analysis (**Figure S3**; **Table S7**), the strongest predictor contributing to the total variation was silt (1.38%), followed by mean temperature of the wettest quarter (0.70%) and precipitation of the coldest quarter (0.55%) (**Figure S4a**; **Table S8**). In comparison, the weakest variable was organic carbon stock in the soil (0.39%).

Environmental variation, geographical distance, and population processes can strongly influence the detection of adaptive signals. In our redundancy analysis (RDA), we partitioned the environmental variables, geographical distances, and population structure to show the contribution of each in explaining the observed genomic variation (**Table 1**). In a full RDA, environmental factors, geographic distance, and population structure explained 16.1 % (R2 adj = 0.120; p=0.000) of the total genetic variation. Further partitioning showed that environmental factors explained 5.0% (R2 adj = 0.022; p=0.042) of the SNP variation, while geographical distance did not have a significant effect on the SNP variation explained (R2 adj = 0.002; p=0.061), suggesting an association between genetic variation and the environmental gradient. These results indicate that environmental factors are stronger predictors of genetic variation than geographic distance among sample collection sites. These results were comparable to other studies in wild currant tomatoes (Gibson and Moyle, 2020) and wild barley (Chang *et al*., 2022).

**Table 1:**
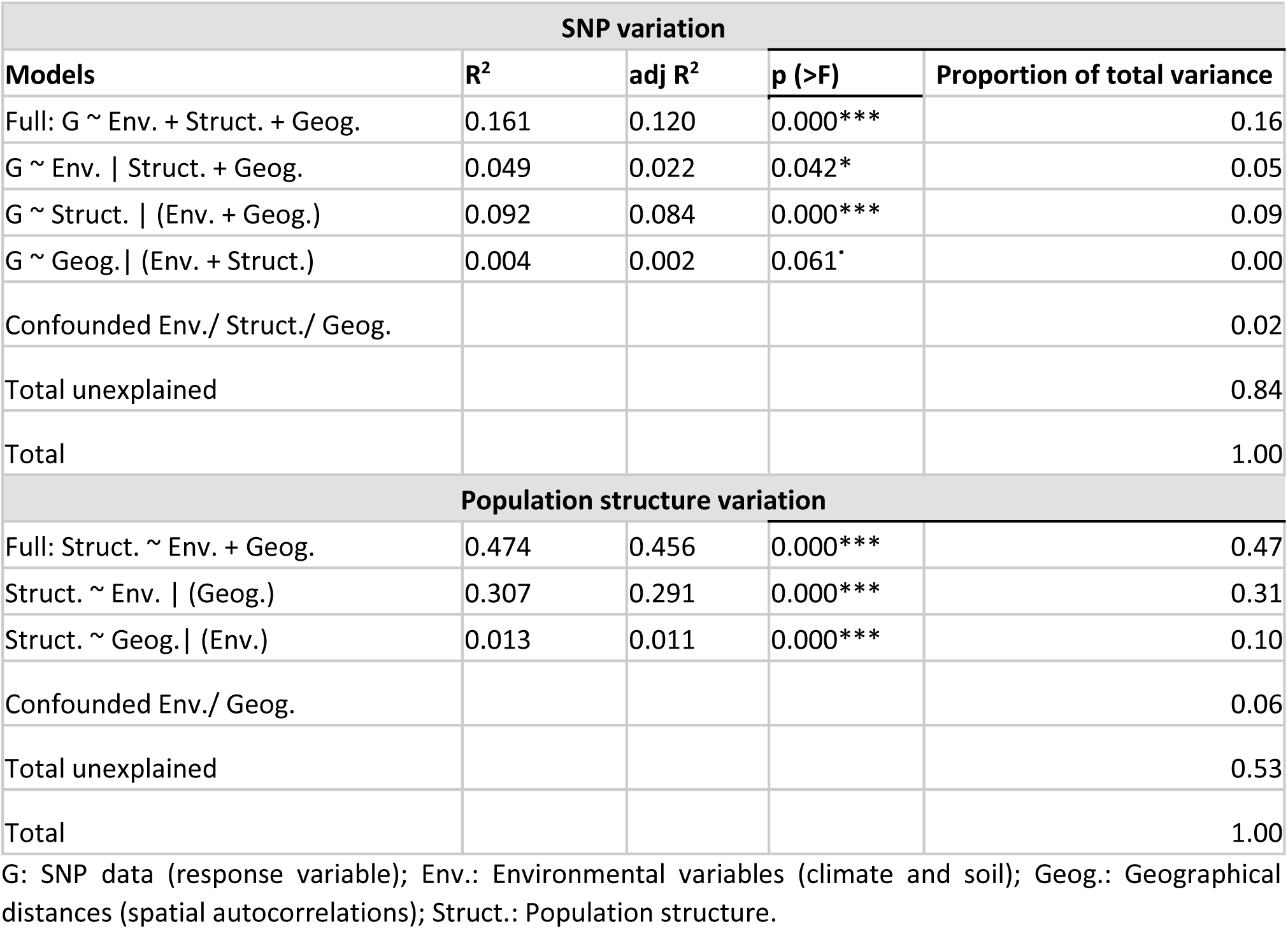
The contribution of the environmental variables (climatic and soil), population structure, and geographical distances to the genomic variation and population structure in the partial RDA models.

Climate plays a vital role in shaping the genetic variation of different crops (Lasky *et al*., 2012; Abebe *et al*., 2015; McGaughran *et al*., 2014). Studies in other crops have examined population structure as a factor besides geography and environment (Capblancq and Forester, 2021; Chang *et al*., 2022). In our case, population structure only explained 9% (R2 adj = 0.084; p=0.000). As is the case in many landscape genomics studies (Chang *et al*., 2022; Wang *et al*., 2023), a large proportion of the genetic variation remained unexplained. In our study, there are two possible reasons for this phenomenon. Firstly, despite our comprehensive analysis of numerous environmental factors, additional unmeasured ecological dynamics, such as biotic interactions and agricultural practices, may be at play. While biotic factors may be partially associated with the abiotic factors examined in this study, as indicated by the observation that various potential adaptive genes are associated with biotic interactions (**Table S9**), agricultural practices such as artificial irrigation, fertilization, pest control treatments, and weed management are more difficult to capture. Lastly, RDA can only establish linear associations between the environmental factors/geographical distances (Borcard *et al*., 2018) and, therefore, may fail to account for any potential non-linear correlations that might be present.

We also wanted to understand how much of the population structure was influenced by environmental and spatial factors, given that the genetic clusters aligned with eco-geographical habitats. Environmental factors and geographic distances accounted for 47% (R2 adj = 0.456; p=0.000) of the population structure (**Table 1**). The environment accounted for most of the population structure at 31% (R2 adj = 0.291; p=0.000). Geographic distances accounted for only 10% (R2 adj = 0.011; p=0.000). This result confirms that the environments of the populations in the Indian subcontinent significantly influence the population structure. However, geography still makes a significant contribution. Demography and geography do not always differentiate when partitioned, since isolation by distance is usually the main driver for intraspecific genetic structure (Lasky *et al*., 2015).

### Main environmental factors contributing to genomic divergence

Our findings identified environmental factors that uniquely contribute to the genomic variation in the Indian subcontinent. After controlling geographic distance and population structure, the RDA biplots show the correlation between genetic variation and environmental predictors (**Figure S4b; Table S10**). The highest biplot scores on the first axis (28.6%) were silt (0.691) and mean temperature of the driest quarter (−0.375). The highest on the second and third axes were for precipitation of the coldest quarter (−0.619 and-0.621, respectively). The correlations between the environmental variables and the RDA axes revealed the effect of the environmental variables on the population variations depending on the geographical locations across the sampling area. This emphasizes the vital role of the environment in facilitating divergent selection among the populations in our study. For instance, populations from south India (Cls_3) experience higher precipitation in the coldest quarter; the southern and central India (Cls_3 and Cls_4) populations experience higher temperatures during the driest season. Central India (Cls_4) populations grow on soils with significantly lower nitrogen content and organic carbon (**Figure S5**). The significant differences in the climatic and soil features experienced by the four population clusters may contribute to the divergent selection observed in the regional clustering.

Our findings on the individual effects of the environmental variables on the SNP variation in both simple and partial RDA, when conditioned on population structure, showed that only silt content in the soil, precipitation of the coldest quarter, precipitation of the driest month, and mean temperature of the driest quarter were significantly associated (p<0.05) with the SNP variation (**Figure S4a; Table S10**). The results suggest that variation in precipitation and soil texture may represent the main drivers of genomic variation. There was a minimal decrease in the explained SNP variation by silt, precipitation of the driest month, and precipitation of the coldest quarter when the population structure was controlled (**Figure S4a**), indicating that these variables correlated less with the population structure compared to mean temperature in driest quarter. This is further supported by the ANOVA analysis of the environmental characteristics of the population clusters (**Figure S5**).

### Detection of adaptive candidate SNPs by different methods

The combination of different methods to identify potential adaptive SNP can reduce false positive rates (Martins *et al*., 2018; Lu *et al*., 2019). The four methods applied detected 305 outlier SNPs in total (305/4308-7.1%, q-value <= 0.05) (**Figure S6; Table S9**) while the sRDA, pRDA, LFMM, and PCAdapt identified 110, 54, 62, and 111 candidate SNPs, respectively (**Figure 5; Figure S6; Figure S7**). Overall, 29 SNPs of the total candidate SNPs were commonly detected by at least two of the methods, with a majority (22 SNPs) by partial RDA and LFMM (**Table S11**). Two SNPs were shared between PCAdapt and LFMM, four between simple RDA and LFMM, and one between simple RDA and PCAdapt. Limited overlaps of SNPs among methods is expected due to the different working background assumptions of the methods, as shown by previous studies applying similar methods (Chang *et al*., 2022). Most of the common outlier SNPs are associated with precipitation in the coldest quarter (n=20), nitrogen (n=4), silt (n=3), and one each for the soil cation exchange capacity, precipitation seasonality, mean temperature of the driest month and precipitation of the driest month.

**Figure 5:**
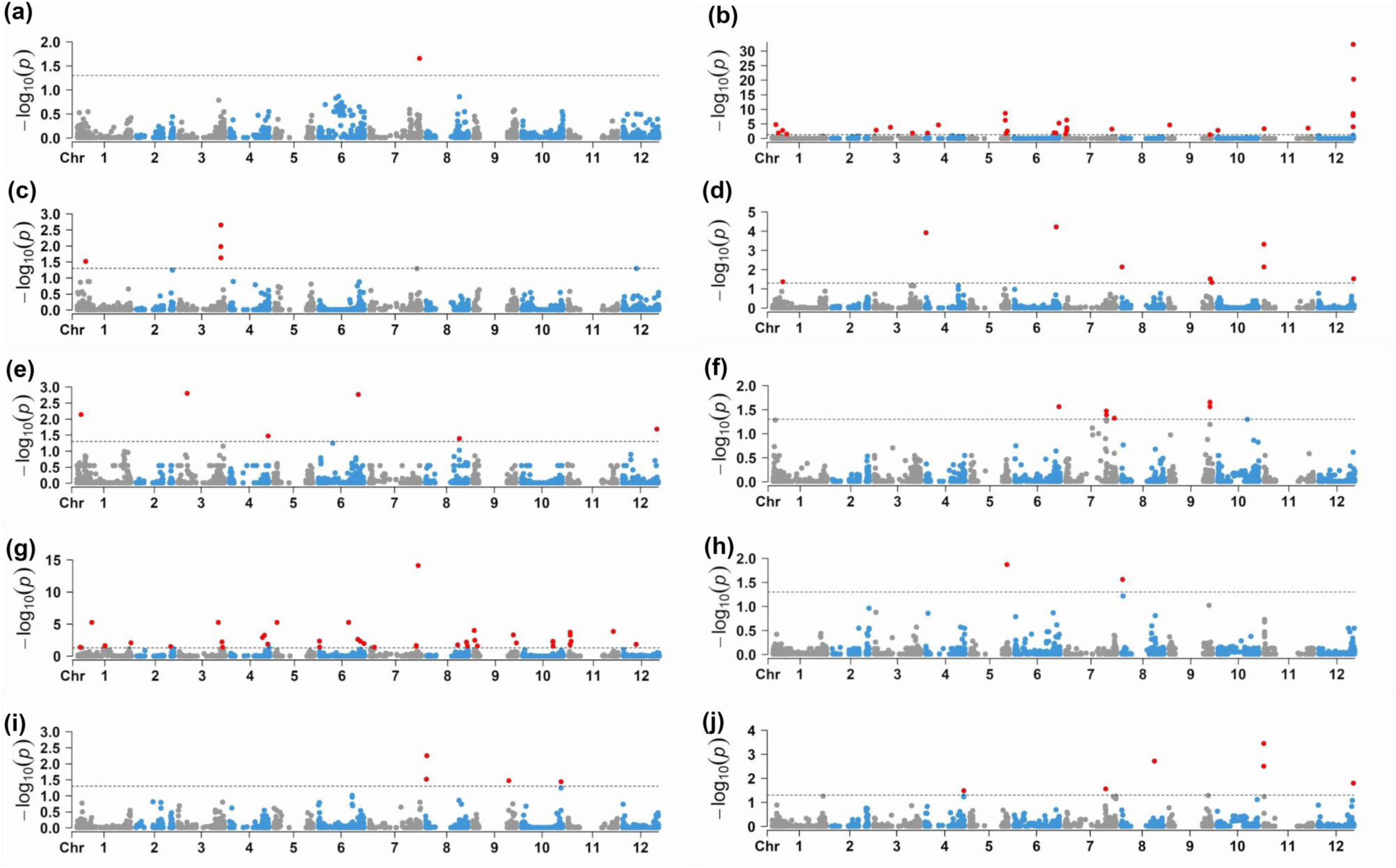
Genome scans for adaptation signatures using LFMM. The Manhattan plots correspond to the 10 environmental variables. Significant SNPs are highlighted as red dots (FDR = 0.05). (a)cec: cation exchange capacity, (b)PCoQ: Precipitation of the coldest quarter, (c)clay: Soil clay content, (d)PDrM: Precipitation of the driest month, (e)Silt: Soil silt content, (f)Pse: Precipitation seasonality, (g)nitro: Soil nitrogen content, (h)PWaQ: Precipitation of the warmest quarter, (i)ocs: Soil organic carbon stock, (j)MTDrQ: Mean temperature of the driest quarter.

### Candidate genes associated with local environmental adaptation

We associated all 305 identified candidate SNPs (**Table S9**) detected by RDA, LFMM, or PCAdapt with documented genes and proteins using the eggplant genome version 4.1 (Barchi *et al*., 2021). These genes were identified from previously identified orthologs in *Arabidopsis thaliana* (https://www.arabidopsis.org/) and other species, including *Nicotiana tabacum, N. benthamiana, Pisum sativum, Vitis vinifera, Citrus sinensis, Mentha piperita, Oryza sativa, and Solanum lycopersicum*.

Our interpretation of the data primarily centers around shared SNPs found within genes related to stimuli response or other genes relevant to the environment (**Figure S6; Table S11**). By focusing on these specific SNPs, we could better understand the climatic factors that influence the current patterns of adaptive variation, as well as the function of the associated genes, facilitating the mining of adaptive loci for breeding purposes. Our study effectively identifies interesting genes showing direct involvement in adaptive pathways related to abiotic stress. Precipitation of the coldest quarter (PCoQ_19), silt content in the soil, and mean temperature of the driest quarter (MTDrQ_9) were the environmental factors associated with most of the genes.

We identified several genes involved in stress hormone pathways, e.g., *SCE1* (SUMO-conjugating enzyme). SCE1 is essential in mediating the conjugation of a small ubiquitin-like modifier (SUMO) to target proteins.

Increased SUMO conjugates in plants are associated with increased abscisic acid (ABA) or abiotic stresses such as cold, high salinity, and heat (Nurdiani *et al*., 2018; Augustine *et al*., 2016; Castro *et al*., 2012; Chaikam and Karlson, 2010). ABA is associated with reactions to drought stress, such as stomatal closure. Also, SUMO machinery components tend to be strongly expressed during seed development in maize, indicating its implication in seed survival under normal and stress conditions (Augustine *et al*., 2016). We also identified *AZI1* (*AZELAIC ACID INDUCED1*), usually induced following pathogen infection and whose overexpression has also been observed to lead to improved freezing tolerance in Arabidopsis (Atkinson *et al*., 2013; Xu *et al*., 2011).

We identified several regulatory genes with potential roles in plant stress responses (e.g., *At1g03560*, *BSK7*, *LRK10*). *AtLRK10L1.2*, the Arabidopsis ortholog of wheat *LRK10*, is involved in ABA-mediated signaling and drought resistance (Lim *et al*., 2015), and also controls flowering time and defense responses against pathogens (Allen *et al*., 2007). *At1g03560* is part of a large pentatricopeptide repeat protein family, playing a role in organelle development through their binding to organellar transcripts (Lurin *et al*., 2004). *BSK7* is a probable serine/threonine kinase that acts as a positive regulator of brassinosteroids-steroidal hormones regulating multiple physiological and developmental processes in plants, including flowering induction and stress responses (Sreeramulu *et al*., 2013; Bajguz and Hayat, 2009; Li *et al*., 2010).

Several identified genes were associated with different aspects of reproductive development and general plant growth, such as regulating pollen tube growth, controlling flowering time, regulating the transition from vegetative to reproductive phase, leaf and root structure development, and cell division. These genes included *CPK17*, encoding a calcium-dependent protein kinase that is essential for pollen fitness (Liu *et al*., 2023; Chen *et al*., 2021); *PPC* (p*hosphoenolpyruvate carboxylase*) essential for growth, development, and seed quality (Feria *et al*., 2022)*; RLT1* is a regulator protein required for the maintenance of the plant vegetative phase (Li *et al*., 2012); *AZI1* (*AZELAIC ACID INDUCED 1*) – a lipid transferase protein that seems to control flowering and lignin synthesis besides conferring cold tolerance (Shi *et al*., 2011; Xu *et al*., 2011); *RID1* (*root initiation defective1-1*) has been shown to be temperature sensitive for adventitious and lateral root formation (Ohtani *et al*., 2013)*; CEL1* is an endoglucanase that plays a role in cell wall relaxation during cell growth and expansion (Tsabary *et al*., 2003); *Dynamin-related protein 3A* (*DRP3A*) is highly expressed in flower tissues and is involved in peroxisome and mitochondria fission (Lingard *et al*., 2008). *REVOLUTA* (*REV)* encodes a transcription factor regulating the relative growth of apical and non-apical meristems (Tsabary *et al*., 2003). The onset of reproduction is a crucial stage in a plant’s life cycle, and natural selection can optimize the most appropriate time and conditions for flowering to ensure reproductive success (Sinha *et al*., 2022).

Intracellular trafficking pathways are crucial for the regulation of stress response mechanisms. They significantly impact essential cellular processes by aiding the movement of diverse molecules and ions across the plasma membrane. This movement is necessary for maintaining ion homeostasis, adjusting osmotic balance, facilitating signal transduction, and aiding detoxification (Vishwakarma *et al*., 2019). We identified a suite of genes associated with transport across membranes. *Vascular protein sorting 18* (*VPS18*) involved in pathways to the protein storage vacuoles (Rojo *et al*., 2003) are upregulated in abiotic stress conditions, including hydric stress (Neves *et al*., 2021). This gene was associated with precipitation of the coldest quarter. Other transport membrane proteins included *At4g32640* (Qu *et al*., 2014) and *At3g30340* (Busov *et al*., 2004).

A few candidate genes appeared to be involved in plant pathogen resistance. *RIN1* (Holt *et al*., 2002), *PUB26* negatively regulates immunity by marking BKI kinases whose phosphorylation and accumulation are central to immune signal propagation (Wang *et al*., 2018). These pathogen resistance genes were associated with precipitation of the coldest quarter. The dispersal and infection success of phytopathogens is greatly enhanced by precipitation and abundant moisture (Milici *et al*., 2020). This may indicate that, in addition to climate, other factors, including pathogens, insects, and herbivores, could contribute to the diversity and local adaptation of the eggplant populations.

We also detected a gene *PU1* (Wattebled *et al*., 2008) involved in starch metabolism. Starch metabolism resulting in sugar accumulation has been described in the literature as a mechanism for drought response, possibly because sugar accumulation helps the plant to stabilize its membranes and proteins to resist dehydration (Perdiguero *et al*., 2012; Pommerrenig *et al*., 2018). Lastly, six genes did not have any known function. These genes are interesting candidates for future investigations to unravel the molecular basis of fitness in different environmental conditions.

### Evidence for distinct genomic signatures of anthropic vs. environmental selection in the Indian domestication center

Approximately 12,000 ago, with the onset of agriculture, humans started selecting plants with desirable characteristics for cultivation (Milla, 2023; Diamond, 2002). However, during early agriculture, humans had much lower control on the agroecosystems than they exert today with irrigation, fertilizers and crop protection agents. It is therefore likely that, besides human selection, local environmental factors present in the domestication centers such as climate, geography, pathogens, soil composition, exerted selective pressures on early cultivars, directly affecting the adaptive traits and associated regions in the genome (Zhao *et al*., 2013).

In order to discriminate anthropic from environmental selection in the Indian domestication center, we compared the five more significant QTLs identified in the GWA (*corcol_a*, *frucol_a*, *frucur*, *frushr* and *fruape*) with the five more significant regions associated with environmental selection (PDrM_14, PCoQ_19, MTDrQ_9, nitro, and silt), searching for overlapping (anthropic and environmental) vs exclusive (anthropic or environmental) QTLs (**Figure 6**). Of the 133 QTLs analyzed in this approach, 115 were exclusive, considering the linkage disequilibrium of the Indian subcontinent accessions (**Table S12**), while 18 anthropic QTLs overlapped or were located at less than 1Mbp from environmental ones (**Table S13**).

**Figure 6:**
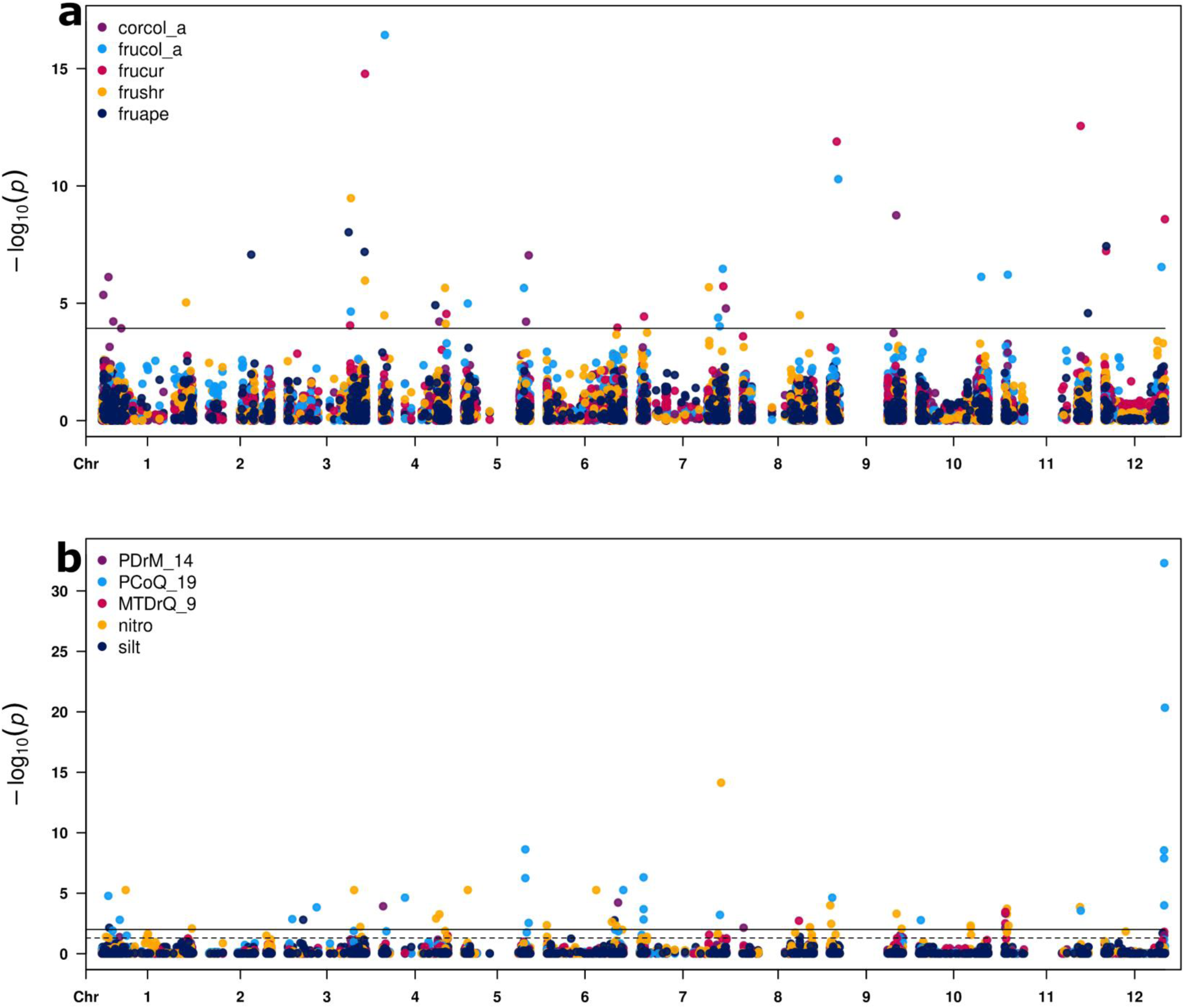
Comparison of the effects of anthropic and environmental selection on the eggplant genome in the Indian domestication center. The 5 QTNs showing the most significant associations by GWA and the 5 adaptive candidate SNPs showing the most significant associations by GEA are shown, respectively, in (a) and (b).

Among the overlapping QTLs, the ones for *frushr* and nitro on chromosome 4 contains a gene coding for IRE (Inositol-Requiring Enzyme), while the ones for silt and *frushr* on chromosome 8 contain an *APC5: Anaphase-promoting complex subunit 5* gene, encoding a subunit of a complex, APC/C, highly conserved among eukaryotes, which plays a key role during gametogenesis, growth, hormone signaling, symbiotic interactions, and endoreduplication onset (Heyman and De Veylder, 2012). Altering *SlCCS52A* (an activator of APC/C) expression altered endoreduplication and fruit size in tomato, in keeping with the *frushr* trait associated with this QTL (Mathieu-Rivet *et al*. 2010)

The QTL for *frucol_a* on chromosome 7 contains a BHLH117, a member of the basic helix-loop-helix (bHLH) transcription factors. The anthocyanin biosynthetic genes are transcriptionally regulated by a MYB-bHLH-WD40 (MBW) complex (Xu *et al*., 2015). In the vicinity, a *PEX22* gene was identified in the QTL for nitrogen content of the soil. Nitric oxide is an important signaling molecule which influences processes as root growth, photomorphogenesis, the hypersensitive response, programmed cell death, stomatal closure, flowering, pollen tube guidance and germination (Nyathi and Baker, 2006). Our results suggest that the anthropic selection for fruit color and the natural selection in response to nitrogen content in the soils acted in the same genomic regions.

The QTL for *frucur* on chromosome 7 contains a WAK2 (Wall Associated Kinase 2) gene, which is implicated in signaling pathways that mediate cell wall integrity and stress responses, as well as in controlling cell expansion, morphogenesis and development. In the vicinity, a *AZI1*: pEARLI1-like lipid transfer protein 1 gene was identified in the QTL for PCoQ_19. This lipid transfer protein seems to control the flowering process and lignin synthesis, as well as to prevent electrolyte leakage during freezing damage (Shi *et al*., 2011; Xu *et al*., 2011). Our results suggest that the anthropic selection for fruit curvature and the adaptation to precipitation of the coldest quarter acted in the same genomic regions.

In conclusion, the limited overlap between anthropic and environmental QTLs is likely due to chance, considering the extent of the linkage disequilibrium in the Indian subcontinent population. This suggests that anthropic and environmental selection act largely independently, and on distinct genomic regions. Further refinement of the phenotypic data collected on accessions originating from a precise geographical area, and of the pedoclimatic and environmental conditions prevalent in that area, will allow a better understanding of this point.

## Conclusions

Our study contributes to elucidating the complex interplay between genetics, human and environmental selection in shaping the extraordinary diversity of eggplant. Using historical genebank characterization data and SPET genotyping data covering the whole genome on 3,449 accessions, significant correlations between various traits and genetic markers were identified, and QTNs and QTLs associated to key agronomic traits were detected. We note that the *MYB1* and *BEAT* genes identified as candidate anthocyanin regulators in this work have also been identified by independent approaches (Docimo *et al*., 2016; Sulli *et al*., 2021), thus providing a nice confirmation of the suitability of genebank-collected phenotypic data for associations studies. By leveraging a comprehensive dataset of georeferenced accessions from the Indian subcontinent and applying advanced genome-wide association methods on environmental factors driving genomic divergence over 300 potential adaptive genes have been identified, underscoring the importance of understanding local adaptation for crop improvement. Further environmental factors representing a multifactorial nature of the environment may be considered in future studies as they develop. Reliable global datasets of environmental data sets are also steadily developing and will significantly improve the outcomes of future studies (Dauphin *et al*., 2023). Our work provides a valuable resource for breeding programs aimed at enhancing resilience and productivity in the face of changing climatic conditions.

### Experimental procedures

#### Germplasm collection for GWAS

A total of 3,449 accessions of eggplant and its wild species were provided by seven genebanks, including the International vegetable genebank at the World Vegetable Center (Taiwan), the Leibniz Institute of Plant Genetics and Crop Plant Research (IPK, Germany), the Germplasm Bank at the Universitat Politècnica de València (UPV-COMAV, Valencia, Spain), the Centre de Ressources Biologiques Légumes de l’Unité de Génétique et Amélioration des Fruits et Légumes (GAFL, INRAE, Montfavet, France), the Centre for Genetic Resources (CGN, Wageningen, The Netherlands), as well as by the genebanks of the Batı Akdeniz Agricultural Research Institute (BATEM, Antalya, Turkey) and the Council for Agricultural Research and Economics (CREA, Montanaso Lombardo, Italy) (see Barchi *et al*., 2023 for their detailed description).

### Genome-wide association (GWA) analysis

We analyzed historical phenotypic data available for twenty-three qualitative/pseudo-qualitative descriptors referring to three trait categories (plant, inflorescence, and fruit) for 438 (fruit size) to 1,519 (fruit color at commercial ripeness) accessions (**Table S1; Table S2**). Different genebanks assessed the phenotypic traits during their characterization trials based on standardized morphological descriptors (ECPGR, 2008; IBPGR, 1990; Taher *et al*., 2017; van der Weerden and Barendse, 2007). Before analysis, all data were reviewed and harmonized, removing any inconsistency (e.g., traits not registered likewise or outliers). For fruit colour at the commercial stage and corolla colour, discontinuous values were converted in tristimulus parameters L*, a*, and b* (representing lightness, redness, and yellowness, respectively) as well as to hexadecimal values (from RGB coordinates, according to the formula: D = ((R * 256) ˄ 2 + (G* 256) + B).

We carried out the GWA analysis using MLMM (Segura *et al*., 2012), FarmCPU (Xiaolei Liu *et al*., 2016) and BLINK (Huang *et al*., 2019) methods implemented in GAPIT3 (Wang and Zhang, 2021), using target SPET SNPs (see Barchi *et al*., 2023) with a MAF >0.05 as molecular markers. The significance threshold for marker-trait associations (MTA) was set at α = 0.01 using the FDR value according to the Benjamini–Hochberg procedure. We generated the Manhattan plots for each trait using the CMplot R package (https://github.com/YinLiLin/R-CMplot). To identify candidate genes, LD regions of significant SNPs were investigated, while overall QTLs were identified by merging overlapping LD regions for the 23 descriptors in the study using bedtools (Quinlan and Hall, 2010). GWA analyses were also carried out separately on the India and Southeast Asian material, and accessions from these regions were identified and analyzed as previously described using BLINK (Huang *et al*., 2019) method.

### Landscape genomics analysis

We georeferenced 324 and 342 accessions from the Indian subcontinent and Southeast Asia, respectively (**Table S14; Table S15; Figure S8**). Among the Southeast Asian accessions, 195 are from Thailand and 147 are from the Philippines. We focused our landscape genomics analysis on the Indian subcontinent domestication center because: i) It presents a diverse pedo-climatic environment that already includes the climates occurring in Thailand and the Philippines, where the vast majority of georeferenced samples from Southeast Asia were collected (**Figure S7**); ii) It allows to assess environmental adaptation across a continuous geographic gradient that allows both geneflow and selection process to happen, in contrast to the geographic isolation between Thailand and the Philippines, that might lead to altered signals of local adaptation and genetic drift due to limited geneflow; iii) fine-grained geoclimatic data (see below) were more readily retrievable for Indian subcontinent compared to Southeast Asia. We categorized the populations from Indian subcontinent into Southern, Central, Eastern, and North/ Northeastern India and those from Bangladesh as Southern, Central, and Northern regions.

We obtained bioclimatic variables related to temperature precipitation, solar radiation, wind speed, and vapor pressure from WorldClim 2.0 (Fick and Hijmans, 2017) with a 2.5 min (∼5km) resolution. The data represent a 30-year average from 1970 to 2000. We averaged the monthly solar radiation, wind, and vapor pressure rasters to obtain annual value rasters from this period. Soil variables included nitrogen, soil organic carbon, organic carbon density, organic carbon stock, cation exchange capacity, pH, clay sand, and silt content. We downloaded the soil data from the SoilGrids database released in 2016 (https://soilgrids.org/) through ISRIC—WDC Soils (Hengl *et al*., 2017) at 250-meter resolution and at a depth of 15-30 cm, approximated from the eggplant root depth. We aggregated the resolution of the soil dataset to match that of the climate data, ensuring they are consistent in both resolution and extent. The *raster* package in R (Hijmans and Van Etten, 2023) was used to average the aggregated soil values using the *resample* and *extent* functions and to extract the individual sampling point values from the georeferenced collection sites of the accessions. We further selected the bioclimatic and soil variables (hereafter called environmental variables) based on Variable Inflation Factors (VIFs) at a threshold of 5 to mitigate the collinearity problem in the landscape genomics analysis approaches applied. We used the following selected environmental variables in our subsequent analyses: precipitation of the driest month (BIO14), precipitation seasonality (BIO15), precipitation of the warmest quarter (BIO18), precipitation of the coldest quarter (BIO19), mean temperature of the driest quarter (BIO9), soil nitrogen, clay, silt contents, soil organic carbon stock and cation exchange capacity of the soil (**Table S7; Figure S4**). A total set of 4,308 SNPs were employed in the analysis.

### Genotype-Environment Association (GEA) analysis

We used three known landscape genomics approaches that complement each other due to their distinct statistical techniques: redundancy analysis (RDA) (Legendre and Legendre, 2012), latent factor mixed model (LFMM) (Frichot *et al*., 2013), and PCAdapt (Privé *et al*., 2020). RDA and LFMM are GEA approaches that test the association between the environmental and geographic variables with the genetic variation within the populations. To account for the effect of geographic distances on SNP variation in the RDA analysis, we used distance-based Moran’s eigenvector maps (dbMEMs) in RDA (Legendre and Legendre, 2012). To account for the effect of population structure on SNP variation in the partial RDA analysis, we used the ancestry coefficients estimated by the STRUCTURE (Evanno *et al*., 2005) program with the optimal K (K = 4) as covariates. PCAdapt is an outlier differentiation method that does not incorporate environmental variables but identifies outlier loci related to population structure. With this approach, we expect to identify additional candidate SNPs that have not yet been captured by the GEA approaches.

The three sets of variables for our RDA analysis included the ten environmental variables, the dbMEM, and the ancestry coefficients (**Table S8)**. We conducted a full RDA to model the effect of all the variables on the SNP variation. To model the effect of each variable on the SNP variation, we used partial RDA conditioned on covariates, i.e., to estimate the proportion of the SNP variation explained by the environmental variables, geographic distances, and population structure. We performed the RDA using the rda function of the R package vegan (Oksanen, 2010). For all RDA models in this study, we carried out 5,000 permutations to test the significance of explanatory variables with the R function *anova.cca*. We plotted the RDA score to estimate the percentage of variance accounted for by each component and the direction of the effect. The plot also showed each vector depicting the variables’ orientation relative to the RDA axis. After that, we performed an outlier analysis of RDA results to determine the SNPs strongly linked with multivariate environmental gradients. We applied an outlier function (Forester *et al*., 2018) to identify SNPs on the RDA loading with a ±3 standard deviation cutoff.

Given the strong correlation between genetic clusters and eco-geographical habitats, we wanted to understand the extent to which population structure could be attributed to the environmental and geographical distances of the sampling locations. To achieve this, we took a different approach by using ancestry coefficients instead of SNPs in our RDA models. This allowed us to exclude recent genetic variations within populations, enabling a more accurate assessment of the contributions made by environments and geographical distances to the population structure. For this analysis, we replaced the SNPs with ancestry coefficients inferred by STRUCTURE, making the new response variables in our RDA models.

To assess the effect of individual environmental variables on SNP variation, we adopted the method by Chang *et al*. (2022). We systematically introduced one environmental variable as the explanatory variable at a time while treating ancestry coefficients as covariates in our RDA models. Due to the correlations between environmental variables, we performed additional permutation tests to determine the specific effects of these variables. This was achieved by including all environmental variables in a single model and setting the parameter ‘by’ to ‘margin’ in *anova.cca*. By doing so, we could test the significance of each environmental variable while accounting for any confounding effects with other environmental variables.

In LFMM analysis, we controlled the population structure using the optimal *K* (*K* = 4), which we initially determined using the STRUCTURE program. We applied the Markov Chain Monte Carlo algorithm for each variable on the LEA package in R (Frichot and François, 2015) for two runs with a burn-in of 3,000 and 6,000 iterations to compute LFMM parameters (|z|-scores) for all loci. We calculated each locus’s false discovery rate (FDR) q value based on the p values in R (R Development core team). We obtained the candidate SNPs under a false discovery rate at α = 0.05.

The R package *PCAdapt* uses a PCA-based approach to simultaneously infer population structure and identify outlier loci related to this structure (Privé *et al*., 2020). We applied a false discovery rate (FDR = 0.05) as the significance level for detecting the outlier loci.

#### Candidate gene annotation

We identified candidate SNP coordinates on candidate genes using the Sol Genomics Network data FTP database for the eggplant genome consortium version 4.1 (Barchi *et al*., 2021) (https://solgenomics.net/organism/Solanum_melongena/genome). The candidate gene search and gene ontology (GO) terms were assigned using the Arabidopsis information resource (tair) (https://www.arabidopsis.org/) and Uniprot (https://www.uniprot.org/) databases. We characterized the genes and their functions, particularly those associated with abiotic stress and relevant to the adaptation of the eggplant.

## Data availability statement

The VCF genotypic datasets generated for this study are accessible at 10.5281/zenodo.12819501

## Acknowledgments

This work has been funded by the European Union under the grant agreements n. 677379 (Linking genetic resources, genomes, and phenotypes of Solanaceous crops (G2P-SOL)) and n. 101094738 (Promoting a Plant Genetic Resource Community for Europe (PRO-GRACE)).

## Author contributions

JP, SL, EP, NS, GLR, RS, LB and GG conceived the study.

LT, MRT, DA, PF, AB, MJD, JS, VL, HFB, RFMB, MB, RF, AB, MvZ, and RS provided materials and harmonized phenotypic data.

LB, GA, LG produced and analyzed SPET data.

LB and LG carried out GWA analyses on historical data

EO and MvZ performed landscape genomics analyses

EO, LB and LT performed search of candidate genes

LB, EO, and GG wrote the manuscript.

All authors critically revised and approved the manuscript.

## Supporting information

**Figure S1:** Inter-trait Spearman correlations assessed in the mapping population. Colored squares show significant correlations at p< 0 01. Calpri: calyx pricliness; colfle: average color of the flesh; corcol: corolla color at anthesis; floear: flowering earliness from sowing; frcodi: fruit color distribution; fruape: fruit apex shape; frucol: fruit color at commercial ripeness; frucur: fruit curvature; frumdp: fruit cross section; frusec: fruit position of the maximum diameter; frushr: fruit shape length/ breadth ratio; frusiz: fruit size; fruyld: fruit yield per plant; grohab: growth habit; lealob: leaf blade lobes; leapri: leaf prikliness.

**Figure S2:** Bayesian clustering results of the STRUCTURE analysis for eggplant accessions from India and Bangladesh. The number of clusters (K) varied from one to ten in ten independent runs. The corresponding ΔK statistics were calculated according to Evanno *et al.,* (2005).

**Figure S3:** Spearman rank correlation between the environmental variables (current climate and soil variables) selected for GEA analysis.

**Figure S4:** a) The contribution of environmental factors in explaining the percent SNP variation in various models. The simple_single and partial_single models estimate individual effects by fitting one environmental variable at a time. The simple_margin and partial_margin models estimate marginal effects by considering all environmental variables simultaneously. The partial_single and partial_margin models are estimated based on partial RDA, considering the influence of population structure. b) Biplot of the RDA conditioned on population structure and geographic distances. The arrows represent correlations of the environmental factors with the RDA axes (more details in **Table S11**). MTDrQ_9: Mean temperature of the driest quarter, PCoQ_19: Precipitation of the coldest quarter, PWaQ_18: Precipitation of the warmest quarter, PDrM_14: Precipitation of the driest month, Pse_15: Precipitation seasonality, clay: Soil clay content, nitro: Soil nitrogen content, ocs: Soil organic carbon stock, Silt: Soil silt content, cec: cation exchange capacity.

**Figure S5:** The variation of environmental (climate and soil) parameters by regional genetic cluster identified by STRUCTURE analysis. Colors correspond to genetic clusters identified in Figure 1. All panels were significant based on ANOVA (P < 1×10^-8^). Tukey HSD post-hoc comparison results are shown as letters above each box plot.

**Figure S6:** Venn diagram showing the number of significant outlier SNPs detected by GEA and OA methods.

**Figure S7:** Genome scans for adaptation signatures using RDA. Two Manhattan plots correspond to the simple RDA (a), and partial RDA (bOmon) conditioned on population structure and geographical distances between sampling points. Significant SNPs are highlighted as red dots (FDR = 0.05).

**Figure S8:** Geographic distribution of the 790 georeferenced accessions (black dots) from the two domestication centers. Color coding refers to the Köppen climate classification (Beck *et al*., 2018). Af: Tropical rainforest; Aw: tropical savannah, dry winter; As: tropical savannah, dry summer; Am: tropical monsoon; BSh: Dry semi-arid hot; Csa: Temperate, dry hot summer; Cwa: Temperate, dry winter, hot summer; Cwb: Temperate, dry winter, warm summer.

**Table S1:** Passport data, traits and geographical origin of the worldwide collection of *S.melongena* accessions used in the study.

**Table S2:** Total number of significant QTNs identified for the phenotypic traits in the study. Traits are scored according to the IPGR/Bioversity descriptors.

**Table S3:** Historical genebank phenotypic data on a worldwide collection of eggplant accessions.

**Table S4:** QTNs identified according to trait and model. Their position on the genome, the associated p-value, minimum allele frequency (MAF) and the adjusted FDR are reported

**Table S5:** Candidate genes subject to anthropic selection identified by GWAS. For each trait, QTL ID, the model(s) for which the QTL was identified, Gene ID, annotation and putative function of the most relevant genes identified within the confidence interval are indicated. Colocalizing QTL are indicated.

**Table S6:** QTNs identified according to trait and geographical region. Their position on the genome, the associated p-value, minimum allele frequency (MAF) and the adjusted FDR are reported.

**Table S7**: Description of the environmental (climate and soil), geographical distance, and STRUCTURE coefficient variables used in the genome-environment Association (GEA) analysis. The acronyms for the Köppen climates are also described

**Table S8:** Individual effects (simple and marginal effects) of the environmental variable in the RDA analysis.

**Table S9:** SNP identity, chromosome number, gene and gene functions of the 305 candidate SNPs detected by the genome-environment Association (GEA) and outlier detection method.

**Table S10:** RDA Loading of the environmental variable to the RDA axes for partial RDA analysis conditioned on population structure and geographical distances

**Table S11:** Gene annotation and associated variables for 29 potentially adaptive SNPs/ genes commonly detected by at least two methods

**Table S12**: Unique QTLs detected by anthropic and environmental GWA

**Table S13:** Common QTLs between anthropic and environmental GWAS with the candidate genes identified.

**Table S14**: Geolocalization and selected environmental variables for the 324 eggplant accessions from the Indian subcontinent used in the genome-environment Association (GEA) analysis

**Table S15**: Geolocalization (see **Figure S8**) and Köppen climates (see **Table S7**) for the 790 eggplant accessions from the Indian subcontinent and Southeast Asia.

## References

Abebe, T.D., Naz, A.A. and Léon, J. (2015) Landscape genomics reveal signatures of local adaptation in barley (Hordeum vulgare L.). Frontiers in Plant Science, 6. Available at: https://www.frontiersin.org/articles/10.3389/fpls.2015.00813 [Accessed November 11, 2022].

Allen, R.S., Li, J., Stahle, M.I., Dubroué, A., Gubler, F. and Millar, A.A. (2007) Genetic analysis reveals functional redundancy and the major target genes of the Arabidopsis miR159 family. Proceedings of the National Academy of Sciences of the United States of America, 104, 16371–6.

Arrones, A., Mangino, G., Alonso, D., et al. (2022) Mutations in the SmAPRR2 transcription factor suppressing chlorophyll pigmentation in the eggplant fruit peel are key drivers of a diversified colour palette. Frontiers in Plant Science, 13. Available at: https://www.frontiersin.org/articles/10.3389/fpls.2022.1025951 [Accessed November 11, 2022].

Atkinson, N.J., Lilley, C.J. and Urwin, P.E. (2013) Identification of Genes Involved in the Response of Arabidopsis to Simultaneous Biotic and Abiotic Stresses. Plant Physiology, 162, 2028–2041.

Augustine, R.C., York, S.L., Rytz, T.C. and Vierstra, R.D. (2016) Defining the SUMO System in Maize: SUMOylation Is Up-Regulated during Endosperm Development and Rapidly Induced by Stress. Plant Physiology, 171, 2191–2210.

Bajguz, A. and Hayat, S. (2009) Effects of brassinosteroids on the plant responses to environmental stresses. Plant Physiology and Biochemistry, 47, 1–8.

Barchi, L., Acquadro, A., Alonso, D., et al. (2019) Single Primer Enrichment Technology (SPET) for High-Throughput Genotyping in Tomato and Eggplant Germplasm. Front. Plant Sci., 10. Available at: https://www.frontiersin.org/articles/10.3389/fpls.2019.01005/full [Accessed December 12, 2020].

Barchi, L., Aprea, G., Rabanus-Wallace, M.T., et al. (2023) Analysis of >3400 worldwide eggplant accessions reveals two independent domestication events and multiple migration-diversification routes. The Plant Journal, 116, 1667–1680.

Barchi, L., Pietrella, M., Venturini, L., et al. (2019) A chromosome-anchored eggplant genome sequence reveals key events in Solanaceae evolution. Scientific Reports, 9, 11769.

Barchi, L., Rabanus-Wallace, M.T., Prohens, J., et al. (2021) Improved genome assembly and pan-genome provide key insights into eggplant domestication and breeding. The Plant Journal, 107, 579–596.

Beck, H.E., Zimmermann, N.E., McVicar, T.R., Vergopolan, N., Berg, A. and Wood, E.F. (2018) Present and future Köppen-Geiger climate classification maps at 1-km resolution. Sci Data, 5, 180214.

Borcard, D., Gillet, F. and Legendre, P. (2018) Spatial Analysis of Ecological Data. In D. Borcard, F. Gillet, and P. Legendre, eds. Numerical Ecology with R. Use R! Cham: Springer International Publishing, pp. 299–367. Available at: 10.1007/978-3-319-71404-2_7 [Accessed December 13, 2023].

Busov, V.B., Johannes, E., Whetten, R.W., Sederoff, R.R., Spiker, S.L., Lanz-Garcia, C. and Goldfarb, B. (2004) An auxin-inducible gene from loblolly pine (Pinus taeda L.) is differentially expressed in mature and juvenile-phase shoots and encodes a putative transmembrane protein. Planta, 218, 916–927.

Capblancq, T. and Forester, B.R. (2021) Redundancy analysis: A Swiss Army Knife for landscape genomics. Methods in Ecology and Evolution, 12, 2298–2309.

Castro, P.H., Tavares, R.M., Bejarano, E.R. and Azevedo, H. (2012) SUMO, a heavyweight player in plant abiotic stress responses. Cell. Mol. Life Sci., 69, 3269–3283.

CGIAR (2020) Scope and roles of the CGIAR genebanks: 2030 vision. Available at: https://www.nature.com/articles/s43016-020-0074-1 [Accessed July 18, 2024].

Chaikam, V. and Karlson, D.T. (2010) Comparison of structure, function and regulation of plant cold shock domain proteins to bacterial and animal cold shock domain proteins. BMB Reports, 43, 1–8.

Chang, C.-W., Fridman, E., Mascher, M., Himmelbach, A. and Schmid, K. (2022) Physical geography, isolation by distance and environmental variables shape genomic variation of wild barley (Hordeum vulgare L. ssp. spontaneum) in the Southern Levant. Heredity, 128, 107–119.

Chen, X., Ding, Y., Yang, Y., Song, C., Wang, B., Yang, S., Guo, Y. and Gong, Z. (2021) Protein kinases in plant responses to drought, salt, and cold stress. Journal of Integrative Plant Biology, 63, 53–78.

Costa-Neto, G., Crespo-Herrera, L., Fradgley, N., Gardner, K., Bentley, A.R., Dreisigacker, S., Fritsche-Neto, R., Montesinos-López, O.A. and Crossa, J. (2023) Envirome-wide associations enhance multi-year genome-based prediction of historical wheat breeding data. G3 Genes|Genomes|Genetics, 13, jkac313.

Daunay, M.C., Lester, R.N. and Ano, G. (2001) Cultivated eggplants. Tropical Plant Breeding, 200–225.

Dauphin, B., Rellstab, C., Wüest, R.O., Karger, D.N., Holderegger, R., Gugerli, F. and Manel, S. (2023) Re-thinking the environment in landscape genomics. Trends in Ecology & Evolution, 38, 261–274.

Diamond, J. (2002) Evolution, consequences and future of plant and animal domestication. Nature, 418, 700–707.

Docimo, T., Francese, G., Ruggiero, A., et al. (2016) Phenylpropanoids Accumulation in Eggplant Fruit: Characterization of Biosynthetic Genes and Regulation by a MYB Transcription Factor. Front. Plant Sci., 6. Available at: https://www.frontiersin.org/journals/plant-science/articles/10.3389/fpls.2015.01233/full [Accessed November 25, 2024].

ECPGR (2008) Minimum descriptors for eggplant, capsicum (sweet and hot pepper) and tomato.

Evanno, G., Regnaut, S. and Goudet, J. (2005) Detecting the number of clusters of individuals using the software structure: a simulation study. Molecular Ecology, 14, 2611–2620.

FAO (2022) http://faostat3.fao.org/home/E.org/.

Feria, A.B., Ruíz-Ballesta, I., Baena, G., Ruíz-López, N., Echevarría, C. and Vidal, J. (2022) Phosphoenolpyruvate carboxylase and phosphoenolpyruvate carboxylase kinase isoenzymes play an important role in the filling and quality of Arabidopsis thaliana seed. Plant Physiology and Biochemistry, 190, 70–80.

Fick, S.E. and Hijmans, R.J. (2017) WorldClim 2: new 1-km spatial resolution climate surfaces for global land areas. International Journal of Climatology, 37, 4302–4315.

Florio, F.E., Gattolin, S., Toppino, L., Bassolino, L., Fibiani, M., Lo Scalzo, R. and Rotino, G.L. (2021) A SmelAAT Acyltransferase Variant Causes a Major Difference in Eggplant (Solanum melongena L.) Peel Anthocyanin Composition. Int J Mol Sci, 22, 9174.

Forester, B.R., Lasky, J.R., Wagner, H.H. and Urban, D.L. (2018) Comparing methods for detecting multilocus adaptation with multivariate genotype–environment associations. Molecular Ecology, 27, 2215–2233.

Francisco, R.M., Regalado, A., Ageorges, A., et al. (2013) ABCC1, an ATP Binding Cassette Protein from Grape Berry, Transports Anthocyanidin 3-O-Glucosides. The Plant Cell, 25, 1840–1854.

Frichot, E. and François, O. (2015) LEA: An R package for landscape and ecological association studies. Methods in Ecology and Evolution, 6, 925–929.

Frichot, E., Schoville, S.D., Bouchard, G. and François, O. (2013) Testing for Associations between Loci and Environmental Gradients Using Latent Factor Mixed Models. Molecular Biology and Evolution, 30, 1687–1699.

Gibson, M.J.S. and Moyle, L.C. (2020) Regional differences in the abiotic environment contribute to genomic divergence within a wild tomato species. Molecular Ecology, 29, 2204–2217.

Godoy, F. de, Bermúdez, L., Lira, B.S., et al. (2013) Galacturonosyltransferase 4 silencing alters pectin composition and carbon partitioning in tomato. Journal of Experimental Botany, 64, 2449–2466.

Gramazio, P., Prohens, J., Plazas, M., Andújar, I., Herraiz, F.J., Castillo, E., Knapp, S., Meyer, R.S. and Vilanova, S. (2014) Location of chlorogenic acid biosynthesis pathway and polyphenol oxidase genes in a new interspecific anchored linkage map of eggplant. BMC Plant Biology, 14, 350.

He, Y., Li, S., Dong, Y., Zhang, X., Li, D., Liu, Y. and Chen, H. (2022) Fine mapping and characterization of the dominant gene SmFTSH10 conferring non-photosensitivity in eggplant (Solanum melongena L.). Theor Appl Genet, 135, 2187–2196.

Hengl, T., Jesus, J.M. de, Heuvelink, G.B.M., et al. (2017) SoilGrids250m: Global gridded soil information based on machine learning. PLOS ONE, 12, e0169748.

Herrero, J., Santika, B., Herrán, A., Erika, P., Sarimana, U., Wendra, F., Sembiring, Z., Asmono, D. and Ritter, E. (2020) Construction of a high density linkage map in Oil Palm using SPET markers. Scientific Reports, 10.

Heyman, J. and De Veylder, L. (2012) The Anaphase-Promoting Complex/Cyclosome in Control of Plant Development. Molecular Plant, 5, 1182–1194.

Hijmans, R.J. and Van Etten, J. (2023) raster: Geographic analysis and modeling with raster data, Available at: http://CRAN.R-project.org/package=raster.

Holt, B.F., Boyes, D.C., Ellerström, M., Siefers, N., Wiig, A., Kauffman, S., Grant, M.R. and Dangl, J.L. (2002) An Evolutionarily Conserved Mediator of Plant Disease Resistance Gene Function Is Required for Normal Arabidopsis Development. Developmental Cell, 2, 807–817.

https://github.com/YinLiLin/R-CMplot CMplot. Available at: https://github.com/YinLiLin/R-CMplot.

Huang, M., Liu, X., Zhou, Y., Summers, R.M. and Zhang, Z. (2019) BLINK: a package for the next level of genome-wide association studies with both individuals and markers in the millions. GigaScience, 8, giy154.

IBPGR (1990) Descriptors for eggplant.

Langridge, P. and Waugh, R. (2019) Harnessing the potential of germplasm collections. Nat Genet, 51, 200–201.

Lasky, J.R., Des Marais, D.L., Mckay, J.K., Richards, J.H., Juenger, T.E. and Keitt, T.H. (2012) Characterizing genomic variation of Arabidopsis thaliana: the roles of geography and climate. Molecular Ecology, 21, 5512–5529.

Lasky, J.R., Upadhyaya, H.D., Ramu, P., et al. (2015) Genome-environment associations in sorghum landraces predict adaptive traits. Science Advances, 1, e1400218.

Legendre, P. and Legendre, L. (2012) Chapter 11-Canonical analysis. In P. Legendre and L. Legendre, eds. Developments in Environmental Modelling. Numerical Ecology. Elsevier, pp. 625–710. Available at: https://www.sciencedirect.com/science/article/pii/B9780444538680500113 [Accessed October 20, 2022].

Lei, L., Poets, A.M., Liu, C., et al. (2019) Environmental Association Identifies Candidates for Tolerance to Low Temperature and Drought. G3 Genes|Genomes|Genetics, 9, 3423–3438.

Li, G., Zhang, J., Li, J., Yang, Z., Huang, H. and Xu, L. (2012) Imitation Switch chromatin remodeling factors and their interacting RINGLET proteins act together in controlling the plant vegetative phase in Arabidopsis. The Plant Journal, 72, 261–270.

Li, J., Li, Y., Chen, S. and An, L. (2010) Involvement of brassinosteroid signals in the floral-induction network of Arabidopsis. Journal of Experimental Botany, 61, 4221–4230.

Li, S., Wang, C., Zhou, X., Liu, D., Liu, C., Luan, J., Qin, Z. and Xin, M. (2020) The curvature of cucumber fruits is associated with spatial variation in auxin accumulation and expression of a YUCCA biosynthesis gene. Hortic Res, 7, 1–12.

Lim, C.W., Yang, S.H., Shin, K.H., Lee, S.C. and Kim, S.H. (2015) The AtLRK10L1.2, Arabidopsis ortholog of wheat LRK10, is involved in ABA-mediated signaling and drought resistance. Plant Cell Rep, 34, 447–455.

Lingard, M.J., Gidda, S.K., Bingham, S., Rothstein, S.J., Mullen, R.T. and Trelease, R.N. (2008) Arabidopsis PEROXIN11c-e, FISSION1b, and DYNAMIN-RELATED PROTEIN3A Cooperate in Cell Cycle–Associated Replication of Peroxisomes. The Plant Cell, 20, 1567–1585.

Liu, M., Wang, C., Xu, Q., et al. (2023) Genome-wide identification of the CPK gene family in wheat (Triticum aestivum L.) and characterization of TaCPK40 associated with seed dormancy and germination. Plant Physiology and Biochemistry, 196, 608–623.

Liu, Xiao, Guo, L.-X., Jin, L.-F., Liu, Y.-Z., Liu, T., Fan, Y.-H. and Peng, S.-A. (2016) Identification and transcript profiles of citrus growth-regulating factor genes involved in the regulation of leaf and fruit development. Mol Biol Rep, 43, 1059–1067.

Liu, Xiaolei, Huang, M., Fan, B., Buckler, E.S. and Zhang, Z. (2016) Iterative Usage of Fixed and Random Effect Models for Powerful and Efficient Genome-Wide Association Studies. PLOS Genetics, 12, e1005767.

Lu, M., Loopstra, C.A. and Krutovsky, K.V. (2019) Detecting the genetic basis of local adaptation in loblolly pine (Pinus taeda L.) using whole exome-wide genotyping and an integrative landscape genomics analysis approach. Ecology and Evolution, 9, 6798–6809.

Lurin, C., Andreés, C., Aubourg, S., et al. (2004) Genome-Wide Analysis of Arabidopsis Pentatricopeptide Repeat Proteins Reveals Their Essential Role in Organelle Biogenesis[W]. The Plant Cell, 16, 2089– 2103.

Manel, S., Poncet, B.N., Legendre, P., Gugerli, F. and Holderegger, R. (2010) Common factors drive adaptive genetic variation at different spatial scales in Arabis alpina. Molecular Ecology, 19, 3824– 3835.

Martins, K., Gugger, P.F., Llanderal-Mendoza, J., González-Rodríguez, A., Fitz-Gibbon, S.T., Zhao, J.-L., Rodríguez-Correa, H., Oyama, K. and Sork, V.L. (2018) Landscape genomics provides evidence of climate-associated genetic variation in Mexican populations of Quercus rugosa. Evolutionary Applications, 11, 1842–1858.

Mathieu-Rivet, E., Gévaudant, F., Sicard, A., et al. (2010) Functional analysis of the anaphase promoting complex activator CCS52A highlights the crucial role of endo-reduplication for fruit growth in tomato: Endo-reduplication and fruit growth in tomato. The Plant Journal, 62, 727–741.

McGaughran, A., Morgan, K. and Sommer, R.J. (2014) Environmental Variables Explain Genetic Structure in a Beetle-Associated Nematode. PLOS ONE, 9, e87317.

Milici, V.R., Dalui, D., Mickley, J.G. and Bagchi, R. (2020) Responses of plant–pathogen interactions to precipitation: Implications for tropical tree richness in a changing world. Journal of Ecology, 108, 1800–1809.

Milla, R. (2023) Phenotypic evolution of agricultural crops. Functional Ecology, 37, 976–988.

Milner, S.G., Jost, M., Taketa, S., et al. (2019) Genebank genomics highlights the diversity of a global barley collection. Nat Genet, 51, 319–326.

Miyatake, K., Saito, T., Nunome, T., Yamaguchi, H., Negoro, S., Ohyama, A., Wu, J., Katayose, Y. and Fukuoka, H. (2020) Fine mapping of a major locus representing the lack of prickles in eggplant revealed the availability of a 0.5-kb insertion/deletion for marker-assisted selection. Breed Sci, 70, 438–448.

Moglia, A., Florio, F.E., Iacopino, S., et al. (2020) Identification of a new R3 MYB type repressor and functional characterization of the members of the MBW transcriptional complex involved in anthocyanin biosynthesis in eggplant (S. melongena L.) S. Aceto, ed. PLOS ONE, 15, e0232986.

Movahed, N., Pastore, C., Cellini, A., et al. (2016) The grapevine VviPrx31 peroxidase as a candidate gene involved in anthocyanin degradation in ripening berries under high temperature. J Plant Res, 129, 513–526.

Mu, Q., Huang, Z., Chakrabarti, M., Illa-Berenguer, E., Liu, X., Wang, Y., Ramos, A. and Knaap, E. van der (2017) Fruit weight is controlled by Cell Size Regulator encoding a novel protein that is expressed in maturing tomato fruits. PLOS Genetics, 13, e1006930.

Neves, J., Sampaio, M., Séneca, A., Pereira, S., Pissarra, J. and Pereira, C. (2021) Abiotic Stress Triggers the Expression of Genes Involved in Protein Storage Vacuole and Exocyst-Mediated Routes. International Journal of Molecular Sciences, 22, 10644.

Nurdiani, D., Widyajayantie, D. and Nugroho, S. (2018) OsSCE1 Encoding SUMO E2-Conjugating Enzyme Involves in Drought Stress Response of Oryza sativa. Rice Science, 25, 73–81.

Nyathi, Y. and Baker, A. (2006) Plant peroxisomes as a source of signalling molecules. Biochimica et Biophysica Acta (BBA)-Molecular Cell Research, 1763, 1478–1495.

Ohtani, M., Demura, T. and Sugiyama, M. (2013) Arabidopsis ROOT INITIATION DEFECTIVE1, a DEAH-Box RNA Helicase Involved in Pre-mRNA Splicing, Is Essential for Plant Development. The Plant Cell, 25, 2056–2069.

Oksanen, J. (2010) Vegan: community ecology package. http://vegan.r-forge.r-project.org/.

Omondi, E.O., Lin, C.-Y., Huang, S.-M., Liao, C.-A., Lin, Y.-P., Oliva, R. and Zonneveld, M. van (2024) Landscape genomics reveals genetic signals of environmental adaptation of African wild eggplants. Ecology and Evolution, 14, e11662.

Ozminkowski, R.H., Gardner, R.G., Henderson, W.R. and Moll, R.H. (1990) Prostrate Growth Habit Enhances Fresh-market Tomato Fruit Yield and Quality. HortScience, 25, 914–915.

Perdiguero, P., Barbero, M.C., Cervera, M.T., Soto, Á. and Collada, C. (2012) Novel conserved segments are associated with differential expression patterns for Pinaceae dehydrins. Planta, 236, 1863– 1874.

Pérez-Díaz, R., Ryngajllo, M., Pérez-Díaz, J., Peña-Cortés, H., Casaretto, J.A., González-Villanueva, E. and Ruiz-Lara, S. (2014) VvMATE1 and VvMATE2 encode putative proanthocyanidin transporters expressed during berry development in Vitis vinifera L. Plant Cell Rep, 33, 1147–1159.

Pommerrenig, B., Ludewig, F., Cvetkovic, J., Trentmann, O., Klemens, P.A.W. and Neuhaus, H.E. (2018) In Concert: Orchestrated Changes in Carbohydrate Homeostasis Are Critical for Plant Abiotic Stress Tolerance. Plant and Cell Physiology, 59, 1290–1299.

Portis, E., Barchi, L., Toppino, L., et al. (2014) QTL mapping in eggplant reveals clusters of yield-related loci and orthology with the tomato genome. PLoS ONE, 9, e89499.

Privé, F., Luu, K., Vilhjálmsson, B.J. and Blum, M.G.B. (2020) Performing Highly Efficient Genome Scans for Local Adaptation with R Package pcadapt Version 4. Molecular Biology and Evolution, 37, 2153– 2154.

Qu, X., Chatty, P.R. and Roeder, A.H.K. (2014) Endomembrane Trafficking Protein SEC24A Regulates Cell Size Patterning in Arabidopsis. Plant Physiology, 166, 1877–1890.

Quinlan, A.R. and Hall, I.M. (2010) BEDTools: a flexible suite of utilities for comparing genomic features. Bioinformatics (Oxford, England), 26, 841–2.

Ranocha, P., Denancé, N., Vanholme, R., et al. (2010) Walls are thin 1 (WAT1), an Arabidopsis homolog of Medicago truncatula NODULIN21, is a tonoplast-localized protein required for secondary wall formation in fibers. The Plant Journal, 63, 469–483.

Rojo, E., Zouhar, J., Kovaleva, V., Hong, S. and Raikhel, N.V. (2003) The AtC–VPS Protein Complex Is Localized to the Tonoplast and the Prevacuolar Compartment in Arabidopsis. MBoC, 14, 361–369.

Sakuraba, Y., Park, S.-Y. and Paek, N.-C. (2015) The Divergent Roles of STAYGREEN (SGR) Homologs in Chlorophyll Degradation. Mol Cells, 38, 390–395.

Satterlee, J.W., Alonso, D., Gramazio, P., et al. (2024) Convergent evolution of plant prickles is driven by repeated gene co-option over deep time, 2024.02.21.581474. Available at: https://www.biorxiv.org/content/10.1101/2024.02.21.581474v1 [Accessed July 22, 2024].

Scaglione, D., Pinosio, S., Marroni, F., et al. (2019) Single primer enrichment technology as a tool for massive genotyping: a benchmark on black poplar and maize. Annals of Botany. Available at: https://academic.oup.com/aob/advance-article/doi/10.1093/aob/mcz054/5424191.

Segura, V., Vilhjálmsson, B.J., Platt, A., Korte, A., Seren, Ü., Long, Q. and Nordborg, M. (2012) An efficient multi-locus mixed model approach for genome-wide association studies in structured populations. Nat Genet, 44, 825–830.

Shi, Y., Zhang, X., Xu, Z.-Y., Li, L., Zhang, C., Schläppi, M. and Xu, Z.-Q. (2011) Influence of EARLI1-like genes on flowering time and lignin synthesis of Arabidopsis thaliana. Plant Biology, 13, 731–739.

Sinha, R., Zandalinas, S.I., Fichman, Y., Sen, S., Zeng, S., Gómez-Cadenas, A., Joshi, T., Fritschi, F.B. and Mittler, R. (2022) Differential regulation of flower transpiration during abiotic stress in annual plants. New Phytologist, 235, 611–629.

Solberg, S., Zonneveld, M. van, Rakha, M., Taher, D., Prohens, J., Jarret, R., Dooijeweert, W. van and Giovannini, P. (2023) Global strategy for the conservation and use of eggplants, Zenodo. Available at: 10.5281/zenodo.7575464.

Song, J., Shang, L., Li, C., et al. (2022) Variation in the fruit development gene POINTED TIP regulates protuberance of tomato fruit tip. Nat Commun, 13, 5940.

Sreeramulu, S., Mostizky, Y., Sunitha, S., et al. (2013) BSKs are partially redundant positive regulators of brassinosteroid signaling in Arabidopsis. The Plant Journal, 74, 905–919.

Stortenbeker, N. and Bemer, M. (2019) The SAUR gene family: the plant’s toolbox for adaptation of growth and development. Journal of Experimental Botany, 70, 17–27.

Sulli, M., Barchi, L., Toppino, L., Diretto, G., Sala, T., Lanteri, S., Rotino, G.L. and Giuliano, G. (2021) An Eggplant Recombinant Inbred Population Allows the Discovery of Metabolic QTLs Controlling Fruit Nutritional Quality. Front. Plant Sci., 12. Available at: https://www.frontiersin.org/articles/10.3389/fpls.2021.638195/full?&utm_source=Email_to_authors_&utm_medium=Email&utm_content=T1_11.5e1_author&utm_campaign=Email_publication&field=&journalName=Frontiers_in_Plant_Science&id=638195 [Accessed May 18, 2021].

Sun, S.-S., Gugger, P.F., Wang, Q.-F. and Chen, J.-M. (2016) Identification of a R2R3-MYB gene regulating anthocyanin biosynthesis and relationships between its variation and flower color difference in lotus (Nelumbo Adans.). PeerJ, 4, e2369.

Taher, D., Solberg, S.Ø., Prohens, J., Chou, Y., Rakha, M. and Wu, T. (2017) World Vegetable Center Eggplant Collection: Origin, Composition, Seed Dissemination and Utilization in Breeding. Frontiers in Plant Science, 8. Available at: https://www.frontiersin.org/articles/10.3389/fpls.2017.01484 [Accessed November 14, 2022].

Tripodi, P., Rabanus-Wallace, M.T., Barchi, L., et al. (2021) Global range expansion history of pepper (Capsicum spp.) revealed by over 10,000 genebank accessions. PNAS, 118. Available at: https://www.pnas.org/content/118/34/e2104315118 [Accessed August 30, 2021].

Tsabary, G., Shani, Z., Roiz, L., Levy, I., Riov, J. and Shoseyov, O. (2003) Abnormal ‘wrinkled’ cell walls and retarded development of transgenic Arabidopsis thaliana plants expressing endo-1,4-β-glucanase (cell) antisense. Plant Mol Biol, 51, 213–224.

Villanueva, G., Rosa-Martínez, E., Şahin, A., García-Fortea, E., Plazas, M., Prohens, J. and Vilanova, S. (2021) Evaluation of advanced backcrosses of eggplant with solanum elaeagnifolium introgressions under low n conditions. Agronomy, 11.

Vishwakarma, K., Mishra, M., Patil, G., et al. (2019) Avenues of the membrane transport system in adaptation of plants to abiotic stresses. Critical Reviews in Biotechnology, 39, 861–883.

Wang, J. and Zhang, Z. (2021) GAPIT Version 3: Boosting Power and Accuracy for Genomic Association and Prediction. Genomics, Proteomics & Bioinformatics. Available at: https://www.sciencedirect.com/science/article/pii/S1672022921001777 [Accessed February 2, 2022].

Wang, Jinlong, Grubb, L.E., Wang, Jiayu, et al. (2018) A Regulatory Module Controlling Homeostasis of a Plant Immune Kinase. Molecular Cell, 69, 493–504.e6.

Wang, Y., Zhang, L., Zhou, Y., Ma, W., Li, M., Guo, P., Feng, L. and Fu, C. (2023) Using landscape genomics to assess local adaptation and genomic vulnerability of a perennial herb Tetrastigma hemsleyanum (Vitaceae) in subtropical China. Front. Genet., 14. Available at: https://www.frontiersin.org/journals/genetics/articles/10.3389/fgene.2023.1150704/full [Accessed July 22, 2024].

Wang, Z., Yang, Z. and Li, F. (2019) Updates on molecular mechanisms in the development of branched trichome in Arabidopsis and nonbranched in cotton. Plant Biotechnology Journal, 17, 1706–1722.

Wattebled, F., Planchot, V., Dong, Y., Szydlowski, N., Pontoire, B., Devin, A., Ball, S. and D’Hulst, C. (2008) Further Evidence for the Mandatory Nature of Polysaccharide Debranching for the Aggregation of Semicrystalline Starch and for Overlapping Functions of Debranching Enzymes in Arabidopsis Leaves. Plant Physiology, 148, 1309–1323.

Weerden, G.M. van der and Barendse, G.W.M. (2007) A web-based searchable database developed for the eggnet project and applied to the radboud university solanaceae database. In Acta Horticulturae. International Society for Horticultural Science (ISHS), Leuven, Belgium, pp. 503–506. Available at: 10.17660/ActaHortic.2007.745.37.

Xiao, X.O., Lin, W. qiu, Li, K., Feng, X.F., Jin, H. and Zou, H. feng (2018) Transcriptome analyses reveal anthocyanin biosynthesis in eggplants, PeerJ Preprints.

Xu, W., Dubos, C. and Lepiniec, L. (2015) Transcriptional control of flavonoid biosynthesis by MYB–bHLH– WDR complexes. Trends in Plant Science, 20, 176–185.

Xu, Z.-Y., Zhang, X., Schläppi, M. and Xu, Z.-Q. (2011) Cold-inducible expression of AZI1 and its function in improvement of freezing tolerance of Arabidopsis thaliana and Saccharomyces cerevisiae. Journal of Plant Physiology, 168, 1576–1587.

Zhang, H., Yu, Z., Yao, X., Chen, J., Chen, X., Zhou, H., Lou, Y., Ming, F. and Jin, Y. (2021) Genome-wide identification and characterization of small auxin-up RNA (SAUR) gene family in plants: evolution and expression profiles during normal growth and stress response. BMC Plant Biology, 21, 4.

Zhang, J., Xu, H., Wang, N., et al. (2018) The ethylene response factor MdERF1B regulates anthocyanin and proanthocyanidin biosynthesis in apple. Plant Mol Biol, 98, 205–218.

Zhang, Y., Hu, Z., Chu, G., Huang, C., Tian, S., Zhao, Z. and Chen, G. (2014) Anthocyanin Accumulation and Molecular Analysis of Anthocyanin Biosynthesis-Associated Genes in Eggplant (*Solanum melongena* L.). Journal of Agricultural and Food Chemistry, 62, 2906–2912.

Zhao, Y., Vrieling, K., Liao, H., et al. (2013) Are habitat fragmentation, local adaptation and isolation-by-distance driving population divergence in wild rice Oryza rufipogon? Molecular Ecology, 22, 5531– 5547.

