## Supplementary figures for "Association analyses reveal both anthropic and environmental selective events during eggplant domestication"

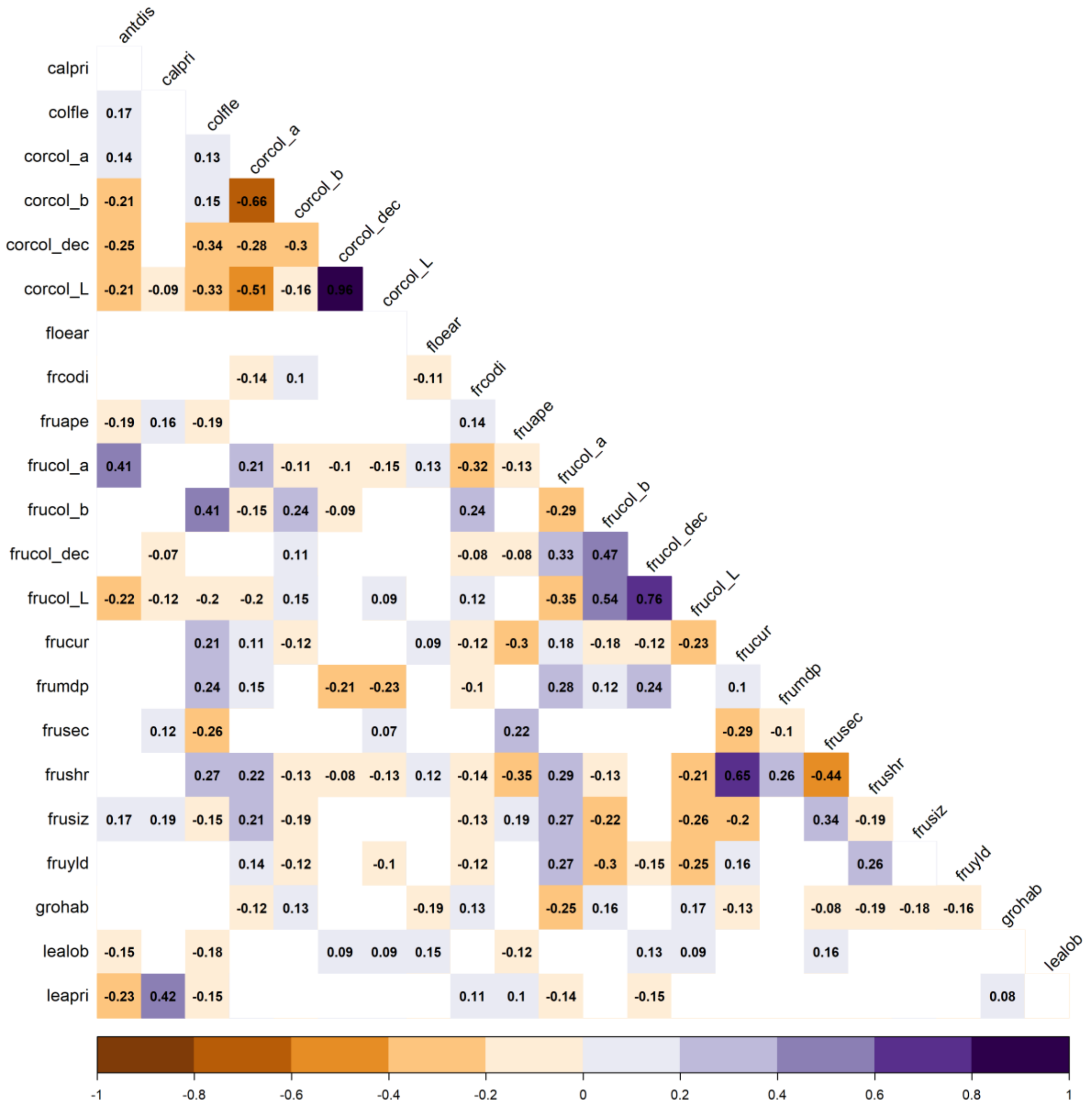


**Figure S1.** Inter-trait Spearman correlations assessed in the mapping population. Colored squares show significant correlations at p< 0 01. Calpri: calyx pricliness; colfle: average color of the flesh; corcol: corolla color at anthesis; floear: flowering earliness from sowing; frcodi: fruit color distribution; fruape: fruit apex shape; frucol: fruit color at commercial ripeness; frucur: fruit curvature; frumdp: fruit cross section; frusec: fruit position of the maximum diameter; frushr: fruit shape length/ breadth ratio; frusiz: fruit size; fruyld: fruit yield per plant; grohab: growth habit; lealob: leaf blade lobes; leapri: leaf prikliness.

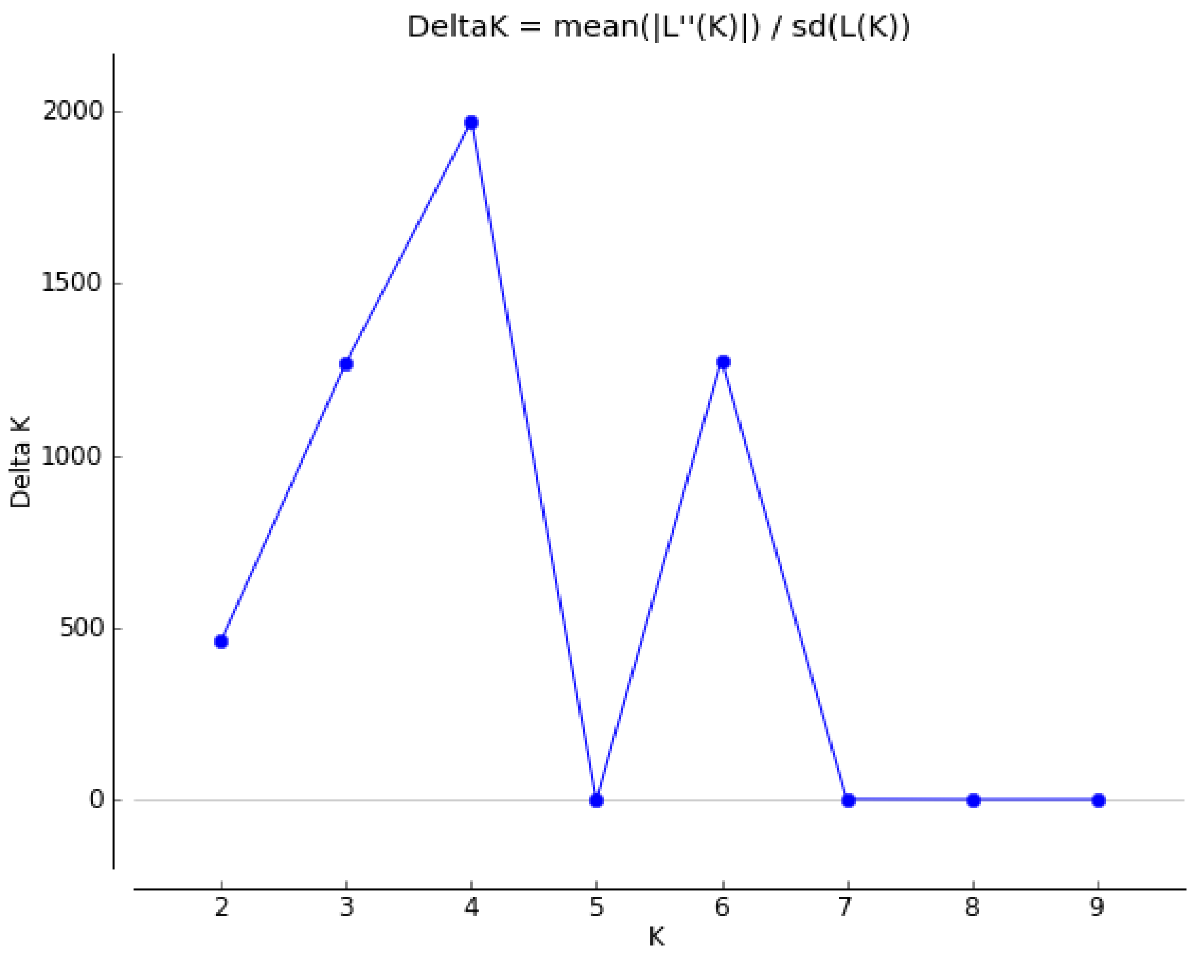


**Figure S2.** Bayesian clustering results of the STRUCTURE analysis for eggplant accessions from India and Bangladesh. The number of clusters (K) varied from one to ten in ten independent runs. The corresponding ΔK statistics were calculated according to Evanno et al. (2005).

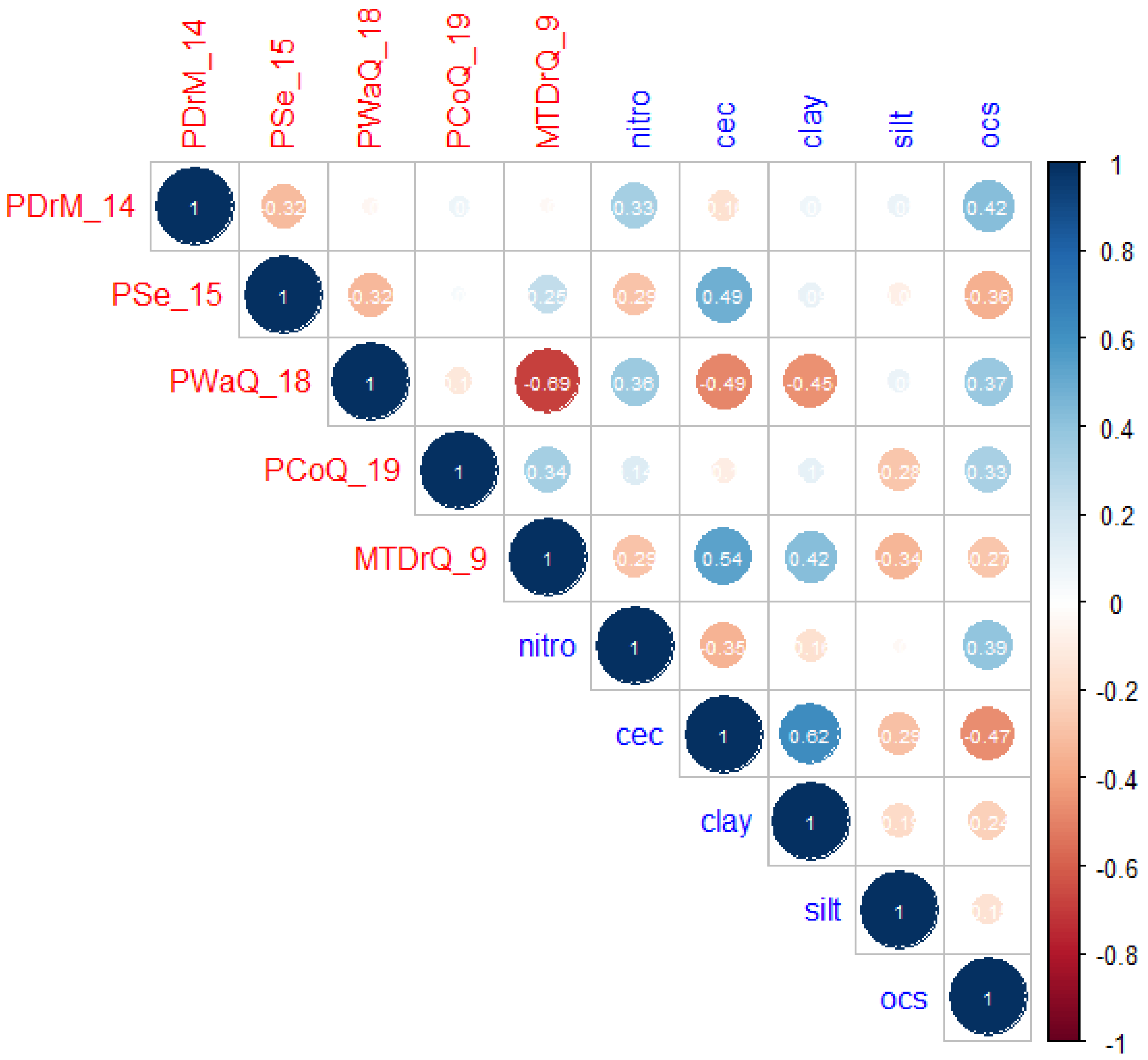


**Figure S3**: Spearman rank correlation between the environmental variables (current climate and soil variables) selected for GEA analysis.

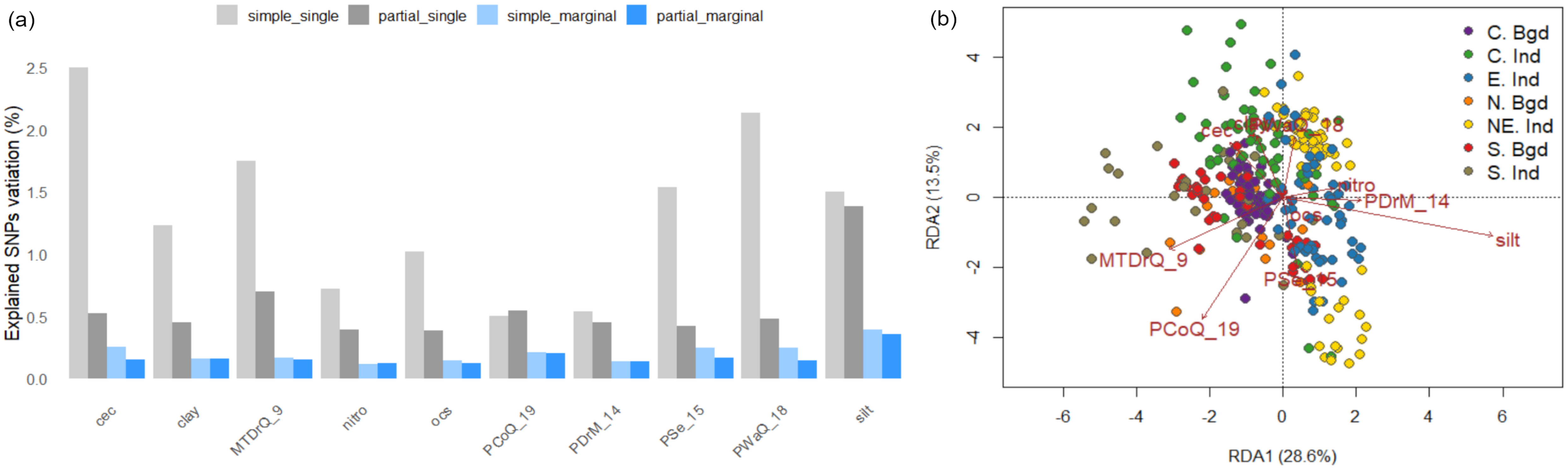


**Figure S4**: **a)** The contribution of environmental factors in explaining the percent SNP variation in various models. The simple_single and partial_single models estimate individual effects by fitting one environmental variable at a time. The simple_margin and partial_margin models estimate marginal effects by considering all environmental variables simultaneously. The partial_single and partial_margin models are estimated based on partial RDA, considering the influence of population structure. **b)** Biplot of the RDA conditioned on population structure and geographic distances. The arrows represent correlations of the environmental factors with the RDA axes (more details in Table S11). MTDrQ_9: Mean temperature of the driest quarter, PCoQ_19: Precipitation of the coldest quarter, PWaQ_18: Precipitation of the warmest quarter, PDrM_14: Precipitation of the driest month, Pse_15: Precipitation seasonality, clay: Soil clay content, nitro: Soil nitrogen content, ocs: Soil organic carbon stock, Silt: Soil silt content, cec: cation exchange capacity.

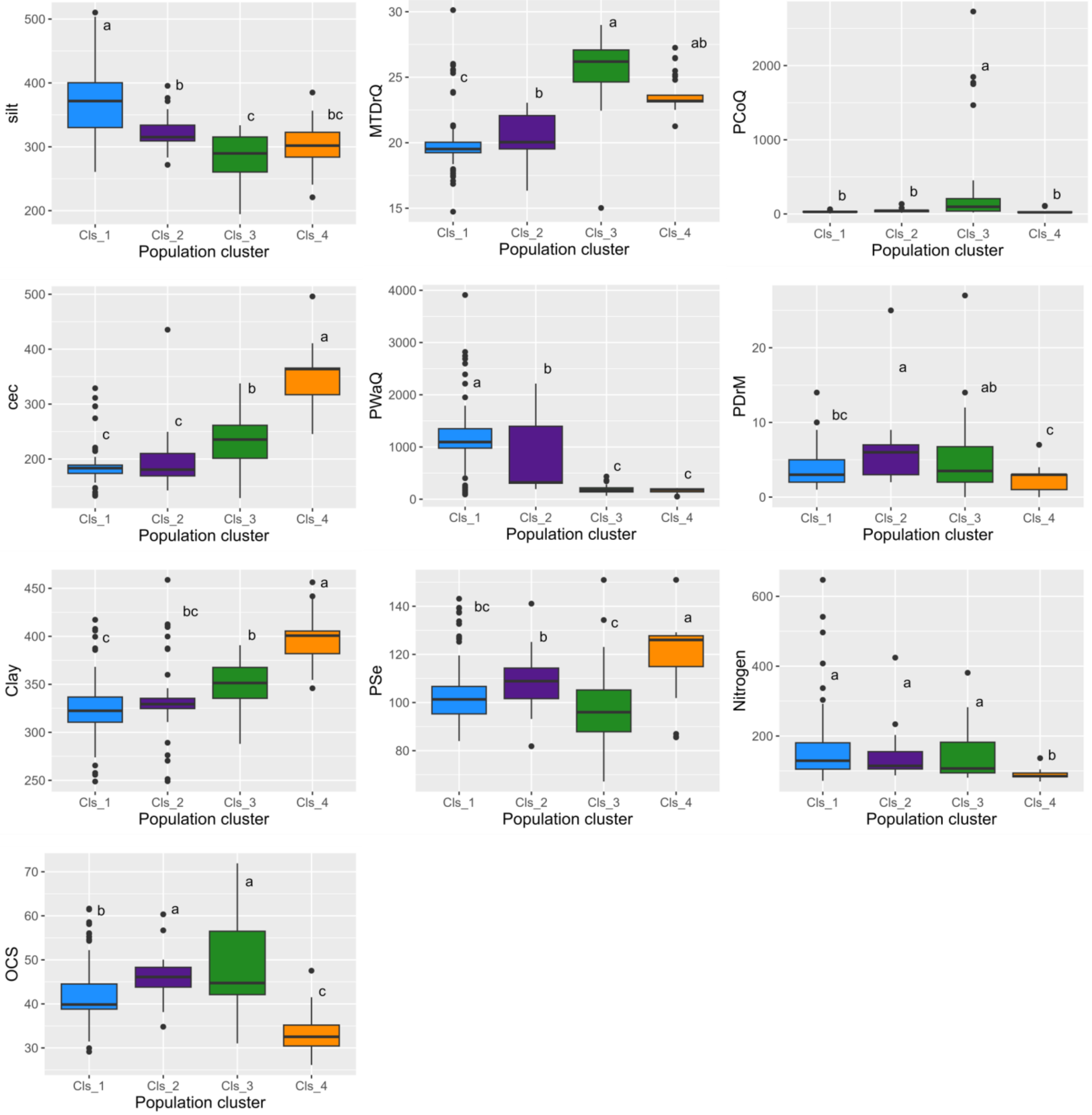


**Figure S5**: The variation of environmental (climate and soil) parameters by regional genetic cluster identified by STRUCTURE analysis. Colors correspond to genetic clusters identified in Figure 1. All panels were significant based on ANOVA ($P < 1\times10^{-8}$). Tukey HSD post-hoc comparison results are shown as letters above each box plot.

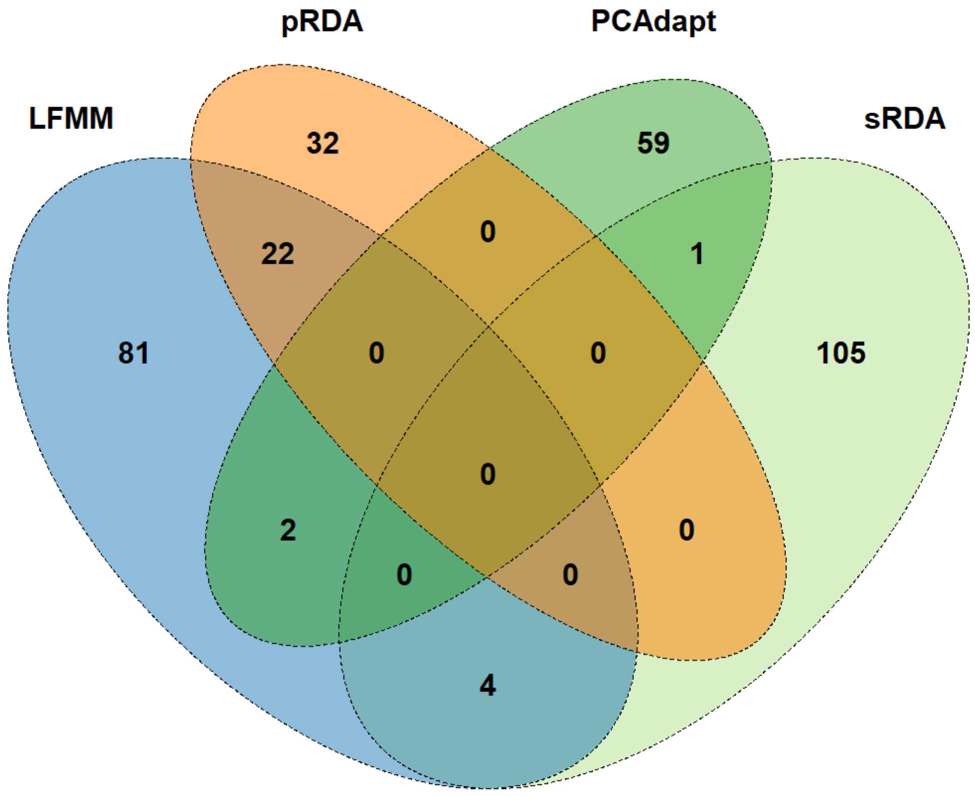


**Figure S6**: Venn diagram showing the number of significant outlier SNPs detected by GEA and OA methods.

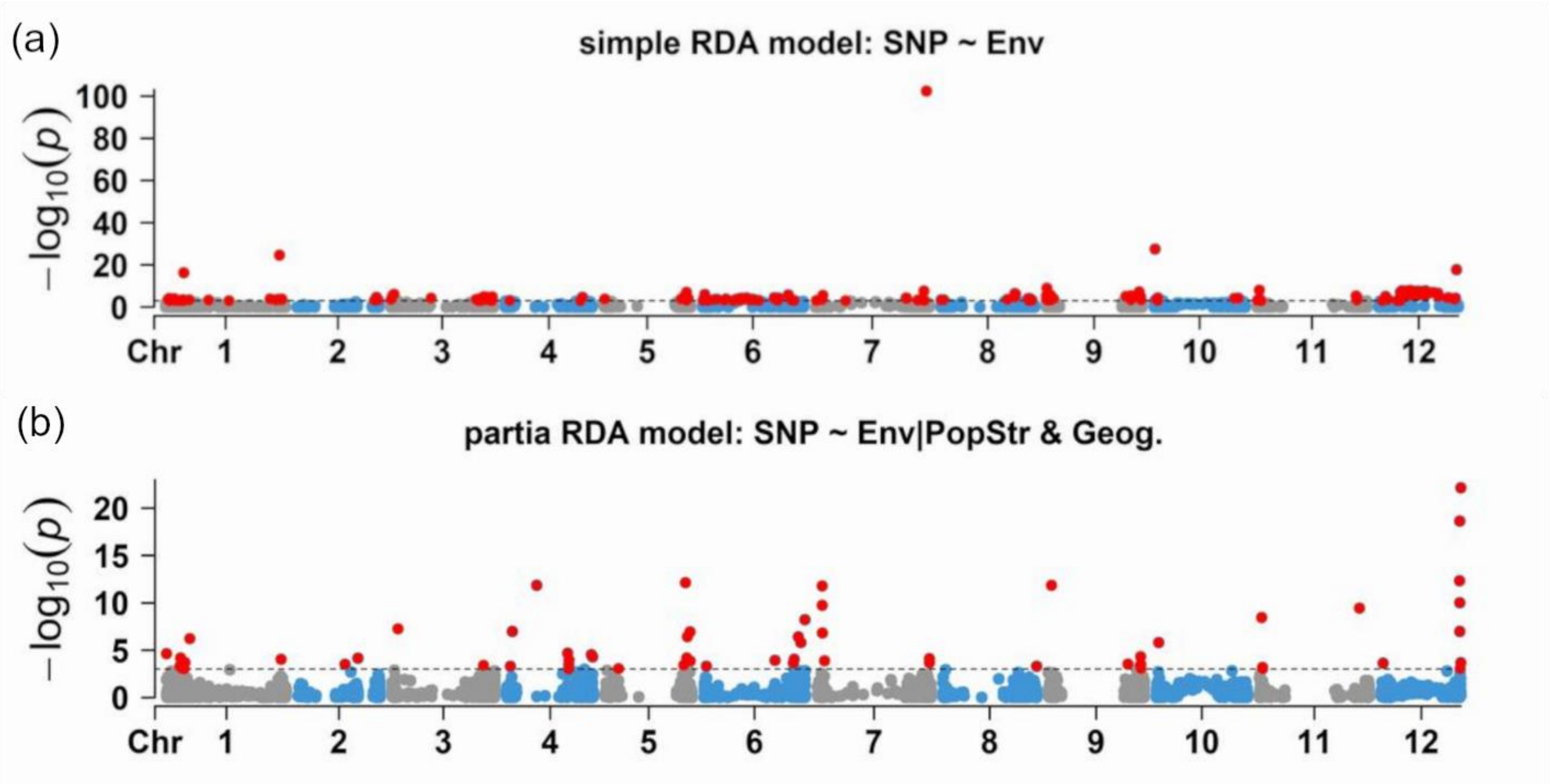


**Figure S7**: Genome scans for adaptation signatures using RDA. Two Manhattan plots correspond to the simple RDA (A), and partial RDA (B) conditioned on population structure and geographical distances between sampling points. Significant SNPs are highlighted as red dots (FDR = 0.05).

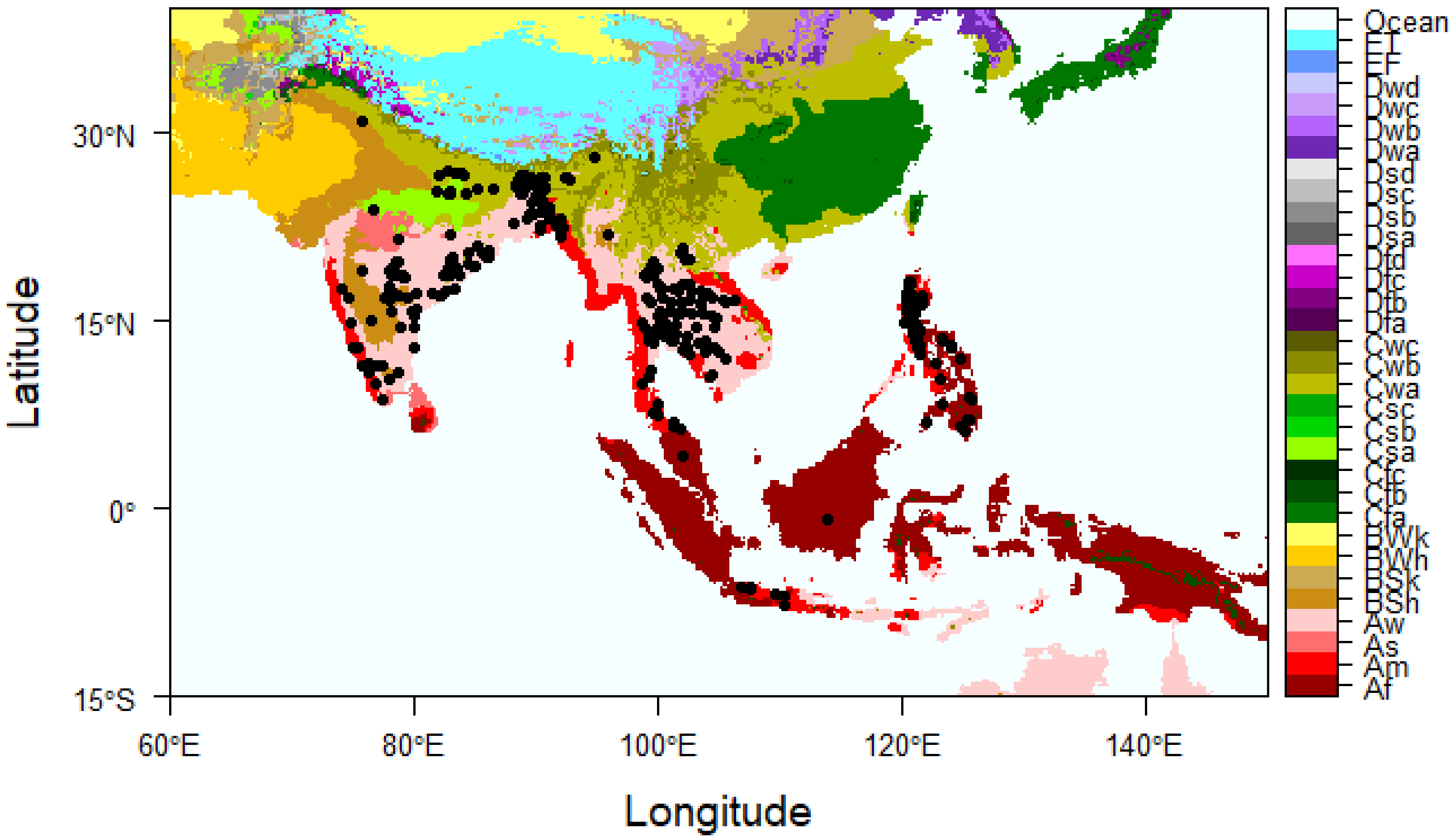


**Figure S8.** Geographic distribution of the 790 georeferenced accessions (black dots) from the two domestication centers. Color coding refers to the Köppen climate classification (https://doi.org/10.1038/sdata.2018.214). Af: Tropical rainforest; Aw: tropical savannah, dry winter; As: tropical savannah, dry summer;  Am: tropical monsoon; BSh: Dry semi-arid hot; Csa: Temperate, dry hot summer; Cwa: Temperate, dry winter, hot summer; Cwb: Temperate, dry winter, warm summer.
